# Molecular mimicry of a pathogen virulence target by a plant immune receptor

**DOI:** 10.1101/2024.07.26.605320

**Authors:** Diana Gómez De La Cruz, Rafał Zdrzałek, Mark J. Banfield, Nicholas J. Talbot, Matthew J. Moscou

## Abstract

Plants and animals respond to pathogen attack by mounting innate immune responses that require intracellular nucleotide binding leucine-rich repeat (NLR) proteins. These immune receptors detect pathogen infection by sensing virulence effector proteins. However, the mechanisms by which receptors evolve new recognition specificities remain poorly understood. Here we report that a plant NLR has evolved the capacity to bind to a pathogen effector by acting as a molecular mimic of a virulence target of the effector, thereby triggering an immune response. The barley NLR Mildew Locus A 3 (MLA3) confers resistance to the blast fungus *Magnaporthe oryzae* by recognizing the effector Pwl2. Using structural analysis, we show that MLA3 has acquired the capacity to bind and respond to Pwl2 through molecular mimicry of the effector host target HIPP43. We demonstrate that the amino acids at the binding interface of MLA3 and Pwl2 are highly conserved in interface of HIPP43 with Pwl2, and are required to trigger an immune response. We used this discovery to bioengineer SR50—an MLA ortholog in rye that confers resistance to wheat stem rust—by introducing the Pwl2 binding interface of MLA3. This chimeric receptor has dual recognition activities, binding and responding to effectors from two major cereal pathogens. Collectively, these results provide evidence that plant immune receptors have evolved sophisticated mimicry strategies to counteract pathogen attack.

## Main Text

Intracellular immune receptors of the nucleotide-binding leucine rich repeat (NLR) class are key components of the innate immune system in eukaryotes. NLRs belong to the superfamily of STAND (signal transduction ATPases with numerous domains) proteins, which are present across the tree of life and oligomerise upon pathogen detection, triggering an inflammatory or cell death response (*1–4*). STAND proteins have a conserved tripartite domain architecture, composed of an N-terminal signalling domain, a central nucleotide-binding NTPase domain and a C-terminal domain that contains superstructure-containing repeats (*5*). In plant NLRs, the central domain is termed NB-ARC (nucleotide-binding adaptor shared by APAF-1, plant R proteins and CED-4) and plays a key role in mediating receptor oligomerisation following pathogen detection (*6*). The N-terminal signalling domain can primarily be of a coiled coil (CC) or Toll/interleukin 1 receptor (TIR) type, and the C-terminal domain comprises leucine rich repeats (LRR), usually associated with pathogen perception (*7, 8*). Broadly, plant NLRs detect pathogen-secreted virulence effectors through direct binding at the LRR domain, or indirectly, by monitoring host proteins modified by effectors (*9, 10*). In some cases, NLR proteins can also contain additional non canonical integrated domains that resemble effector host targets and serve as decoys to activate immunity (*11, 12*). Whether direct or indirect, pathogen perception induces the formation of inflammasome-like NLR oligomers called resistosomes, which initiate immune signalling (*13*). However, the strategies by which new receptor recognition specificities evolve, and the diversity of effector binding interfaces remain mostly unknown.

*Mla3* is a CC type barley NLR that belongs to the *Mildew locus a* (*Mla*) NLR expanded allelic family, known to confer isolate-specific resistance to barley powdery mildew (*Blumeria graminis* f. sp. *hordei* [*Bgh*]), a cereal pathogen of economic significance (*14*). In addition to resistance against *Bgh*, *Mla3* also confers resistance against the blast fungus *Magnaporthe oryzae* by recognising the effector *PWL2* (*Pathogenicity towards weeping lovegrass 2*) (*15, 16*), a widely conserved effector across different host-infecting lineages of *M. oryzae* (*17, 18*). While MLA3 and Pwl2 were previously shown to associate in planta (*15*), the exact recognition mechanism is not yet fully understood. Structural studies of activated plant NLRs upon ligand binding suggest that effectors bind to the ascending lateral chain of the LRR domain to trigger resistosome formation (*19, 20*). Nonetheless, whether this principle applies to all cases of direct NLR-effector binding remains to be determined.

### MLA3 directly binds to Pwl2 through the C-terminus of the LRR domain

Previous work showed that among a series of near-isogenic barley lines with different *Mla* alleles in the Siri genetic background, the accessions S02 and S13 are resistant to *M. oryzae* KEN54-20, unlike their susceptible parent Siri (*Mla8*) (*15*). To confirm that the resistant phenotype in these barley lines is due to recognition of *PWL2* by their *Mla* alleles, we performed spot-inoculation assays with a *M. oryzae* mutant isolate lacking *PWL2*. The near isogenic line S02, which carries *Mla3,* was susceptible to the *pwl2* mutant, indicating specific recognition of *PWL2* in this barley line. S13 showed identical responses to *M. oryzae* as S02 (**Fig. 1A**). Bioinformatic analysis of short and long read transcriptome data indicated that S13 expresses copies of both, *Mla3* and *Mla23* (**Table S1**), suggesting that *PWL2* recognition in this barley line might be conferred either by *Mla3* alone or by both *Mla* alleles. *Mla23,* is the most closely related allele to *Mla3,* with 98.6% protein sequence identity and variation strictly limited to the C-terminus of the NLR (**Fig. 1B**). This prompted us to test whether *Mla23* in S13 also responds and binds to *PWL2* from *M. oryzae.* MLA3 triggers a hypersensitive response when transiently co-expressed with Pwl2 in *Nicotiana benthamiana* and this correlates with association with the effector *in planta* (*15*). To test whether the same is true for MLA23, we transiently co-expressed MLA23 with Pwl2 in *N. benthamiana*, which did not trigger cell death (**Fig. 1C**). In addition, coimmunoprecipitation assays revealed that, unlike MLA3, MLA23 does not associate with Pwl2 (**Fig. 1D**). This indicates that MLA23 does not recognise Pwl2 and resistance to KEN54-20 in S13 is conferred by MLA3 rather than MLA23.

**Fig. 1.**
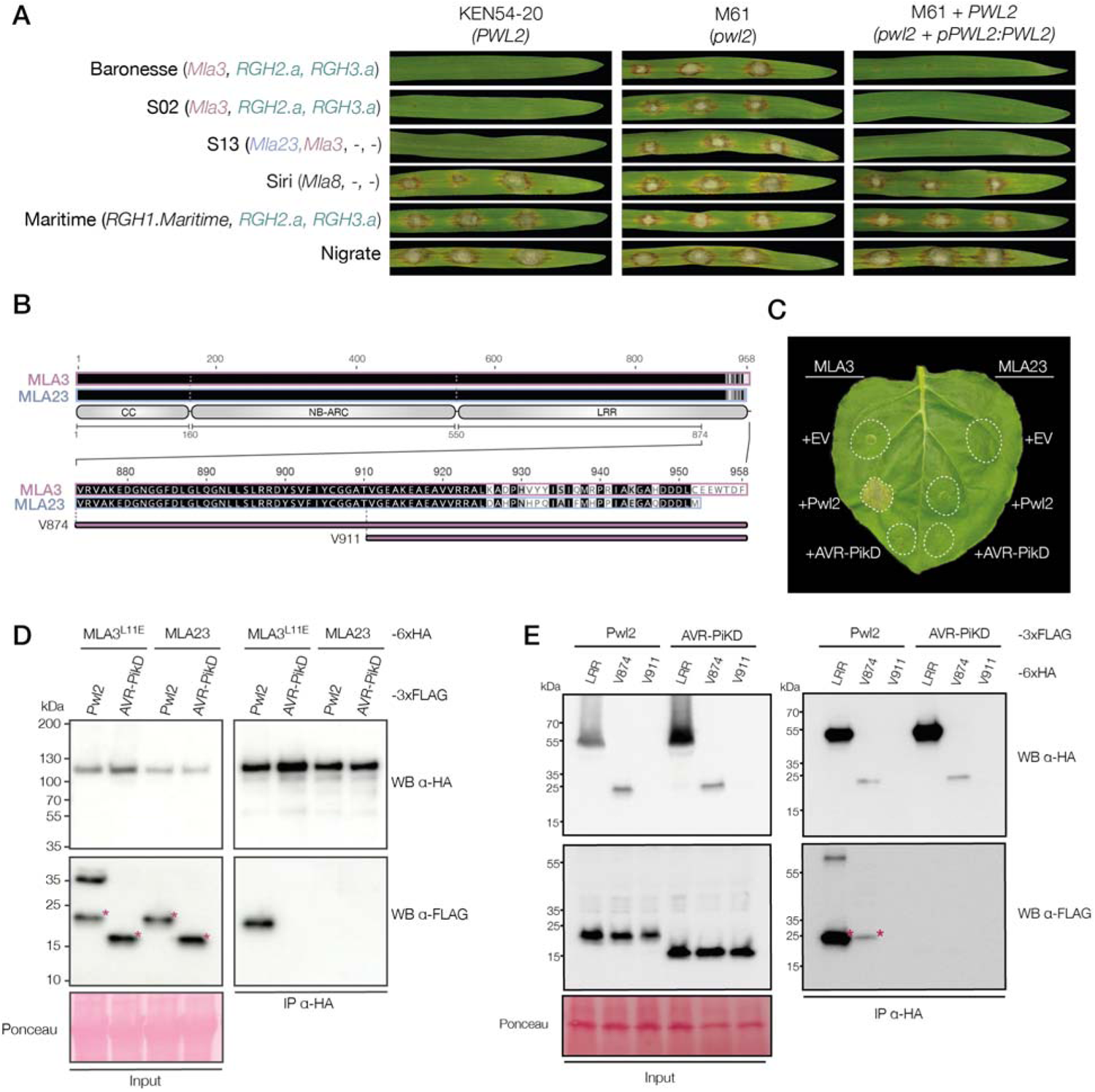
MLA3 recognizes Pwl2 through the C-terminus of its LRR domain. (**A**) Barley leaves spot inoculated with *M. oryzae* KEN54-20 (carrying *PWL2),* a *pwl2* mutant (M61) and a *PWL2*-complemented mutant (M61 + *PWL2*). Each row corresponds to a different barley accession. The *MLA* locus (*RGH1, RGH2* and *RGH3* alleles) of each barley accession is specified in brackets. (**B**) Visual protein alignment of MLA3 vs. MLA23. Black indicates identical residues in both proteins, grey indicates similarity between residues and white highlights lack of conservation. The last 85 amino acids of the alignment are zoomed in to highlight the polymorphisms between proteins. (**C**) Cell death assay in *N. benthamiana* after co-expression of MLA3 or MLA23 with either Pwl2, AVR-PikD or empty vector (EV). The experiment consisted of three biological replicates, each with at least six technical replicates. (**D**) Coimmunoprecipitation assays between MLA3^L11E^ or MLA23 with Pwl2. C-terminally 6xHA-tagged MLA proteins were coexpressed with C-terminally 3xFLAG-tagged fungal effectors. The L11E mutation in MLA3 impairs cell death. AVR-PikD was used as negative control. (**E**) Coimmunoprecipitation assays between LRR fragments of MLA3 shown in (B) and Pwl2. C-terminally 6xHA-tagged MLA3 fragments were coexpressed with C-terminally 3xFLAG-tagged fungal effectors. AVR-PikD was used as negative control. Immunoprecipitations in (D) and (E) were done with agarose beads conjugated to HA antibodies (HA IP). Proteins were detected with the antibodies labelled on the right. Approximate molecular weights of the proteins are shown on the left. Protein loading was checked with Ponceau S solution. Asterisks show the expected protein band for each treatment. The experiments were independently repeated three times with similar results.

MLA23 only differs from MLA3 by 19 amino acid polymorphisms at the C-terminus of the LRR domain (**Fig. 1B**). This suggests that this region of the NLR contains specificity determinants for Pwl2 recognition. To determine which domains of MLA3 are required for association with Pwl2, we generated fragments containing each individual domain of MLA3 (CC, NB-ARC, and LRR), as well as combinations of domains (CC-NB-ARC and NB-ARC-LRR), and tested whether they coimmunoprecipitate with Pwl2. To prevent cell death triggered by activated MLA3, we introduced a mutation in its CC domain (MLA3^L11E^) as this has previously been shown to prevent cell death mediated by activated CC-NLRs (*15, 21*). The NB-ARC-LRR fragment and the LRR domain alone coimmunoprecipitated with Pwl2 (**fig. S1**), indicating that the LRR domain of MLA3 is both necessary and sufficient to bind Pwl2, as expected from previous reports of NLR-effector association (*22–26*).

Given that all amino acid polymorphisms in MLA23 compared to MLA3 are located at the C-terminus of the LRR domain, we hypothesised that this region is crucial for Pwl2 binding. To test if this region is sufficient for Pwl2 binding, we performed coimmunoprecipitation assays with Pwl2 and the MLA3 C-terminal fragments V874 (last 85 amino acids of MLA3) or V911 (last 48 amino acids) (**Fig. 1B**). The MLA3 V874 fragment was sufficient to associate with Pwl2 in *N. benthamiana* protein extracts, whereas the V911 fragment did not accumulate well *in planta* (**Fig. 1D**). Interaction between Pwl2 and the MLA3 V874 fragment was also detected in yeast-two-hybrid assays, indicative of direct binding (**fig. S2**). Overall, these results provide evidence that MLA3 binds to Pwl2 through the C-terminus comprised by the last 85 residues of this NLR.

### MLA3 is predicted to mimic the binding interface of a Pwl2 host target

To better understand the molecular mechanism of Pwl2 recognition by MLA3, we predicted the structure of the LRR domain of MLA3 in complex with Pwl2 using AlphaFold2 (*27, 28*). The best ranked model had pLDDT score = 84.4, pTM score = 0.859, ipTM score = 0.835 and model confidence of 0.839, suggesting high model confidence throughout the structure of the complex, supported by overall low predicted aligned error (**Fig. 2A**). Pwl2 was predicted to interact with the last three C-terminal repeats of the LRR domain of MLA3, which corresponds to the C-terminus of the NLR, in agreement with our experimental data (**Fig. 2A**).

**Fig. 2.**
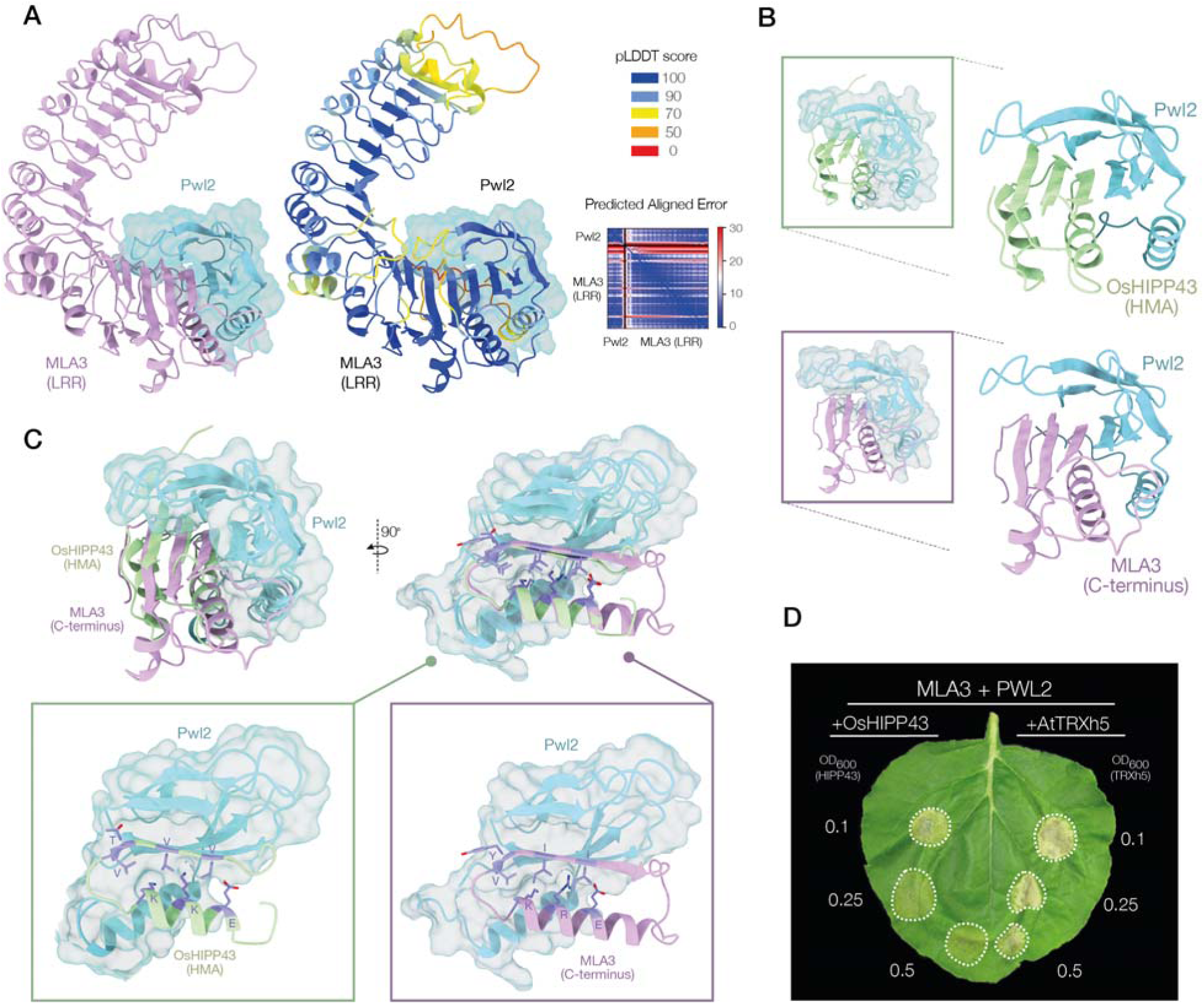
MLA3 mimics the binding interface of a Pwl2 host target. (**A**) AlphaFold2 model of the LRR domain of MLA3 in complex with Pwl2. The structure on the right is colored by pLDDT score, accompanied by the predicted aligned error plot of the model. (**B**) Structure comparison of Pwl2 in complex with its host target OsHIPP43 (PDB ID: 8R7A, upper panel), and the C-terminus of MLA3, according to AlphaFold2 (lower panel). (**C**) Superimposition of the structure of Pwl2 in complex with OsHIPP43 and the model of MLA3 (C-terminus) with Pwl2. Side chains of MLA3 and OsHIPP43 amino acids at the binding interface with Pwl2 are shown for comparison. (**D**) The expression of OsHIPP43 weakens recognition of Pwl2 by MLA3 in *N. benthamiana*. Cell death assay in *N. benthamiana* leaves expressing the indicated constructs. The marked OD_600_ values denote the concentration of the *A. tumefaciens* suspensions used for infiltration carrying OsHIPP43. AtTRXh5 (TRXh5 from *Arabidopsis thaliana*) was used as negative control. MLA3 and Pwl2 were expressed at a constant OD_600_ of 0.5 and 0.4 in all treatments, respectively. The experiment consisted of three biological replicates, each with at least six technical replicates. Protein expression and quantification of cell death phenotypes are shown in fig. S6.

Pwl2 belongs to the structurally conserved family of Magnaporthe Avrs and ToxB-like (MAX) effectors (*29, 30*), and indeed, the structure prediction indicates that Pwl2 has a MAX fold, agreeing with the experimental structure of the effector (*30*) (**fig. S3**). Pwl2 interacts with a Heavy Metal Associated (HMA) protein named HIPP43 as a putative virulence target in rice (OsHIPP43) and barley (HvHIPP43) (*17, 30*), and the structure of the Pwl2-OsHIPP43 complex has been determined (*30*). Structural comparison of the prediction of Pwl2 in complex with the C-terminus of MLA3, and the experimental structure of Pwl2 with OsHIPP43 indicates that Pwl2 binds to both HIPP43 and the C-terminus of MLA3 through a similar, overlapping binding interface (**Fig. 2B**). In addition, in both complexes, MLA3 and OsHIPP43 interact with Pwl2 in related orientations and both interfaces share overall structural features (**Fig. 2B**, **Fig. 2C**), despite the lack of sequence similarity (**fig. S4A**). The HMA domain of OsHIPP43 is formed by four antiparallel β strands and two α-helices, and the C-terminus of MLA3 comprises three parallel β strands and one α-helix (**fig. S4**). These differences in the tertiary structures of HIPP43 and the C-terminus of MLA3 suggest that the region that binds Pwl2 in MLA3 does not correspond to a misannotated integrated HMA domain at the C-terminus of the NLR, but rather corresponds to the last three repeats of the LRR domain.

Even though the order and orientation of the β-strands in both structures is not strictly conserved (**fig. S4**), the α1-helix and subsequent β-strand in each are aligned and oriented in a similar manner to interact with Pwl2 (**Fig. 2C**). We therefore hypothesised that the Pwl2 binding interface of MLA3 shares similarities with the binding interface of OsHIPP43. Analysis with PDBePisa v1.52 (*31, 32*) revealed a total interface area of 1856.1 Å^2^ between MLA3 and Pwl2, compared to 1976.9 Å^2^ in the HIPP43-Pwl2 complex (*30*). The α1-helix and subsequent β-strand in OsHIPP43 and the C-terminus of MLA3 share residues that are identical or have similar properties, and are in direct contact with Pwl2 (**Fig. 2C, fig S5**). In the α1-helix of OsHIPP43, the residues Glu43, Lys47 and Lys51 are buried in the binding interface interacting with Pwl2. Similarly, in the C-terminus of MLA3, the corresponding residues Glu918, Arg922 and Lys926 are precisely located in the same positions at the Pwl2 binding interface. Despite not being identical, both Lys47 from OsHIPP43 and Arg922 from MLA3 are basic, and therefore contribute to comparable physicochemical properties in this area of the binding interface (**Fig. 2C, fig. S5**). In the subsequent β-strand of OsHIPP43, the residues Val56, Thr57, Val59 and Val61 interact with Pwl2. Similarly, in MLA3, the corresponding amino acids Val931, Tyr932, Ile934 and Ile936, located in similar positions in the complex, also interact with Pwl2 (**Fig. 2C, fig. S5**). Valine and isoleucine are amino acids with structurally similar hydrophobic, non-polar side chains. In addition, Thr57 in OsHIPP43 and the corresponding Tyr932 in MLA3 both have a hydroxyl group, conferring analogous features at the binding interface. This model therefore suggests that both plant proteins MLA3 and OsHIPP43 compete to bind Pwl2 through the same interface. To test this, we simultaneously co-expressed MLA3, Pwl2 and OsHIPP43 in *N. benthamiana* and evaluated the impact of OsHIPP43 expression on Pwl2 recognition. Indeed, the cell death response triggered by MLA3 and Pwl2 was affected by the presence of OsHIPP43 (**Fig. 2D, fig. S6**). Overall, these findings suggests that MLA3 has evolved to mimic the binding interface of a Pwl2 host target to detect *M. oryzae* and trigger an immune response.

### MLA3 evolved unique sites at the C-terminus to gain Pwl2 recognition

MLA3 belongs to an extended series of over 30 described alleles that share >90% protein sequence similarity (*14*). All MLA alleles have a structurally conserved C-terminus (**Fig. 3A, fig. S7**). However, only MLA3 is known to confer Pwl2-dependent resistance to *M. oryzae* (*15*). This prompted us to establish whether MLA3 evolved unique sites at the Pwl2 binding interface amongst other MLA alleles that allowed gain of Pwl2 recognition. For this, we performed a phylogenetic analysis of MLA alleles using a codon-based alignment of the full length of the NLR sequences (**Fig. 3A, fig. S8**), extracted the region corresponding to the Pwl2 binding interface in MLA3, and compared the residues in MLA3 predicted to interact with Pwl2 across all MLA alleles (**Fig. 3A, fig. S8**). Most of these amino acids in MLA3 spatially match with HIPP43 residues at the Pwl2 binding interface (**fig. S5**). However, only 3 out of 12 amino acids predicted to bind Pwl2 are completely unique in MLA3 relative to other MLA alleles. These are Lys926, Val931 and Tyr932 (**Fig. 3A, fig. S8**).

**Fig. 3.**
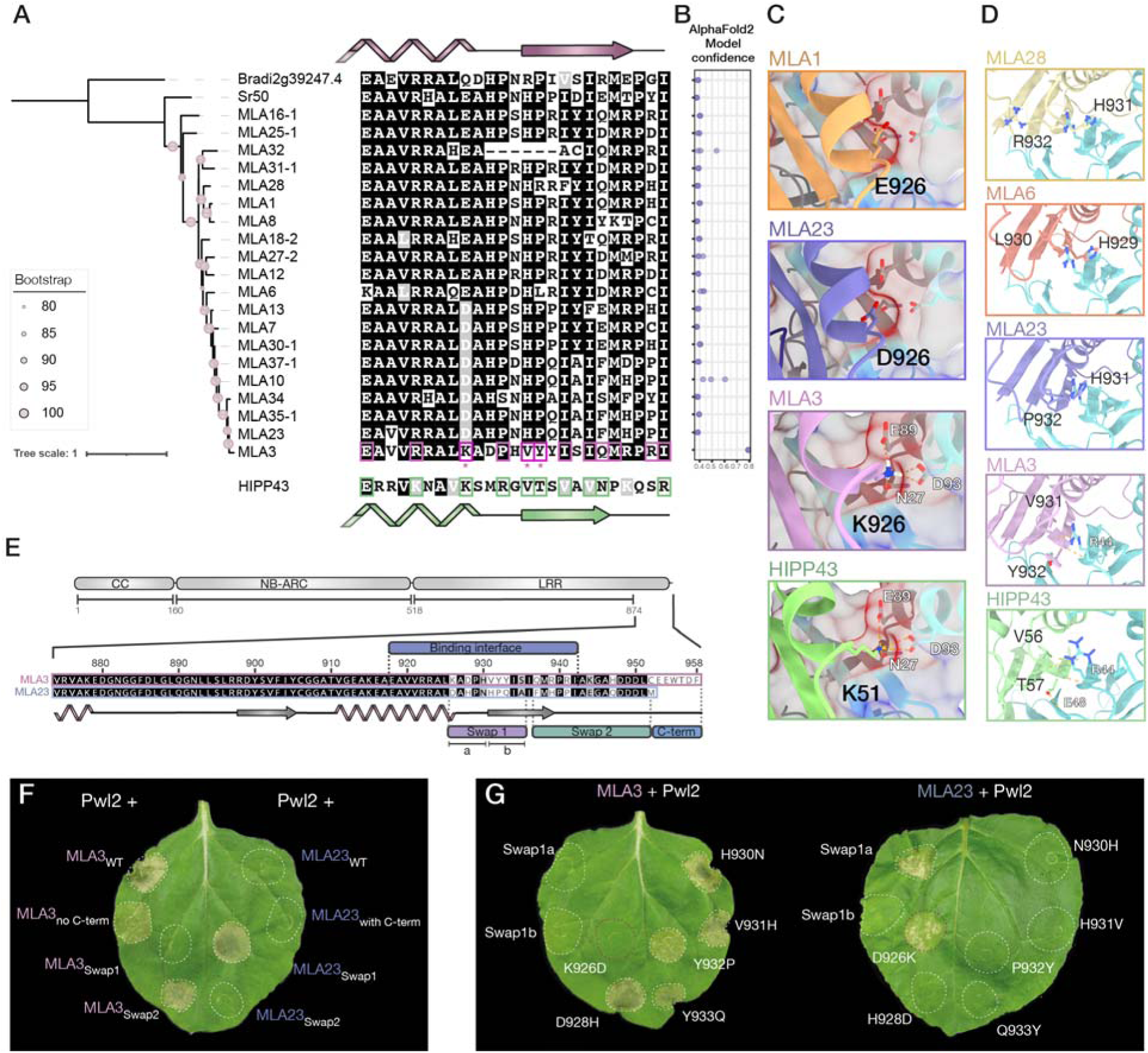
MLA3 has unique sites at the Pwl2 binding interface that allow recognition. (**A**) Phylogenetic tree based on a codon-based alignment of the full sequence of representative MLA alleles and multiple protein sequence alignment of the region corresponding to the Pwl2 binding interface in MLA3. The matching sequence of the binding interface in HIPP43 is shown at the bottom. MLA3 and HIPP43 residues that interact with Pwl2 are highlighted in pink and green boxes, respectively. Amino acids that are unique in MLA3 at the Pwl2 binding interface are shown with an asterisk (K926, V931 and Y932). Schematic secondary structures of the Pwl2 binding interface of MLA3 and HIPP43 are shown at the top and the bottom of the alignment, respectively. (**B**) AlphaFold model confidence score (0.8*ipTM + 0.2*pTM) of the LRR domain of MLA alleles in complex with Pwl2. Structure predictions were performed three independent times. Each dot represents the confidence score of each round of prediction. The analysis with the complete set of MLA alleles is shown in fig. S8. (**C**) Interaction of MLA3^K926^ or HIPP43^K51^ with Pwl2, vs. MLA1^E926^ or MLA23^D926^ as representatives of other MLA alleles. Non-covalent bonds are displayed as dotted yellow lines. (**D**) Interaction of MLA3^V931,Y932^ or HIPP43^V56,T57^ with Pwl2, vs. MLA28^H931,R932^, MLA6^H929,L930^ or MLA23^H930,P931^ as representatives of other MLA alleles. Non-covalent bonds are displayed as dotted yellow lines. (**E**) Schematic representation of the C-terminus of MLA3 and MLA23, highlighting the region corresponding to the Pwl2 binding interface in MLA3, and the regions swapped between MLA3 and MLA23. (**F**) and (**G**) Photos of cell death assays in representative *N. benthamiana* leaves expressing Pwl2 and the indicated chimeric proteins, according to the swaps shown in (E). Reciprocal single amino acid mutations between MLA3 and MLA23 are also shown in (G). The experiments consisted of three biological replicates, each with at least six technical replicates. Protein expression and quantification of cell death phenotypes of (F) and (G) are shown in fig. S10–12.

To test the role of the MLA3 unique sites in the predicted binding interface with Pwl2 compared to other MLA alleles, we predicted the structure of the LRR domain of all known MLA alleles in complex with Pwl2. MLA3 was the only MLA allele predicted by AlphaFold2 to bind Pwl2 with high model confidence (>0.75), averaged from three independent rounds of structure prediction (**Fig. 3B, fig. S8**). Structural alignment of the LRR domain of all MLA alleles with the LRR domain of MLA3 in complex with Pwl2, and assessment of the individual impact of the MLA polymorphisms at the binding interface revealed that MLA3 is the only MLA allele with a basic residue (Lys) at position 926. This lysine corresponds with the Lys51 in OsHIPP43, which forms salt bridges and hydrogen bonds with a negatively charged pocket in Pwl2 made by the amino acids Asn27, Asp93 and Glu89. Lys926 in MLA3 is predicted to interact with the same residues of this pocket in Pwl2. In contrast, all other MLA alleles carry an acidic residue in this position (either Glu or Asp), and due to their negative nature, none of the other MLA alleles are predicted to fit into the negatively charged pocket in Pwl2 where MLA3 Lys926 locates (**Fig. 3C, fig. S9**). In addition, the amino acids Val931 and Tyr932 are unique to MLA3 and match the corresponding OsHIPP43 residues Val56 and Thr57 at the Pwl2 binding interface. In each case, these residues form hydrogen bonds with Arg44 in Pwl2. All other MLA alleles carry His931, followed predominantly by Pro932 in most alleles, Lys in MLA6 or Arg in MLA28. However, the large side chain and the basic nature of His in all other MLA alleles sterically clash with Arg44 in Pwl2 (**Fig. 3D**), likely destabilising the interaction. We suggest that MLA3 evolved unique sites at the binding interface with Pwl2 thereby generating a molecular mimic of the Pwl2 binding interface of HIPP43.

To confirm the role of the key unique residues predicted to be at the Pwl2 binding interface, we evaluated the 19 amino acid differences between MLA3 and MLA23 at the C-terminus by splitting the region comprising all polymorphic sites into three sub-regions: Swap1, Swap2, and the C-terminal extension of MLA3, which is absent in MLA23, but is not predicted to play a role in Pwl2 binding (**Fig. 3E**). We generated reciprocal swaps between MLA3 and MLA23 to test for changes in Pwl2 recognition in cell death assays in *N. benthamiana.* The only MLA3 chimera that completely lost the ability to trigger an immune response upon coexpression with Pwl2 was the one carrying the Swap1 region from MLA23. Conversely, the MLA23 chimera carrying the Swap1 region from MLA3 was the only one that gained the ability to respond to Pwl2 (**Fig. 3F, fig. S10**) This region contains all three unique sites in MLA3 predicted to play an important role in binding of Pwl2 (Lys926, Val931 and Tyr932). To analyse the contribution of each amino acid within the Swap1 region towards Pwl2 recognition, we further split the region into S1a and S1b (**Fig. 3E**), and generated reciprocal chimeras and single amino acid substitutions for each polymorphic site. The mutation D926K in MLA23 was sufficient to confer Pwl2 recognition, and likewise, the reciprocal mutation K926D in MLA3 was sufficient to abolish the cell death response to Pwl2 (**Fig. 3G, fig. S11, fig. S12**). The MLA23^D926K^ mutation was sufficient to result in gain of Pwl2 binding (**fig. S13**), thus confirming the predicted critical role of Lys926 in MLA3. When replaced by all possible amino acids, only the K926R mutation maintained recognition of Pwl2 by MLA3 (**fig. S14**), consistent with the requirement of a positively charged residue in this position to interact with Pwl2. In all cases, absence of cell death was not due to lack of protein expression, as all MLA3 and MLA23 mutants and chimeras accumulated *in planta* (**fig. S10, fig. S11, fig. S14**).

### Bioengineering of SR50 lead to gain of Pwl2 recognition

SR50 is the MLA orthologue in rye and confers resistance against wheat stem rust (*Puccinia graminis* f. sp. *tritici* [*Pgt*]) by recognising the cognate effector AvrSr50 (*33, 34*). SR50 and MLA3 share 77% protein sequence identity, and residues governing specificity of AvrSr50 recognition have been mapped to the ascending lateral chain of the LRR of SR50, up until position 870 (*35*). This prompted us to test whether SR50 can be engineered to gain Pwl2 binding and recognition. For this, we generated variants of SR50 that carry the C-terminus of MLA3 (SR50^3Cterm^), or only the predicted Pwl2 binding interface (SR50^3BI^) (**Fig. 4A**). Both chimeric variants of SR50 respond to Pwl2 when tested in cell death assays in *N. benthamiana* (**Fig. 4B**, **Fig. 4C, fig. S15**). Interestingly, the variant strictly containing the Pwl2 binding interface also retained AvrSr50 recognition (**Fig. 4C, fig. S15**). In both cases, gain of Pwl2 recognition correlated with gain of Pwl2 binding *in planta* (**Fig. 4D**). Therefore, the bioengineered SR50^3BI^ carries two distinct functional binding interfaces to recognise structurally distinct effectors from two major cereal fungal pathogens (**Fig. 4E, fig. S16**) (*17, 30, 36*).

**Fig. 4.**
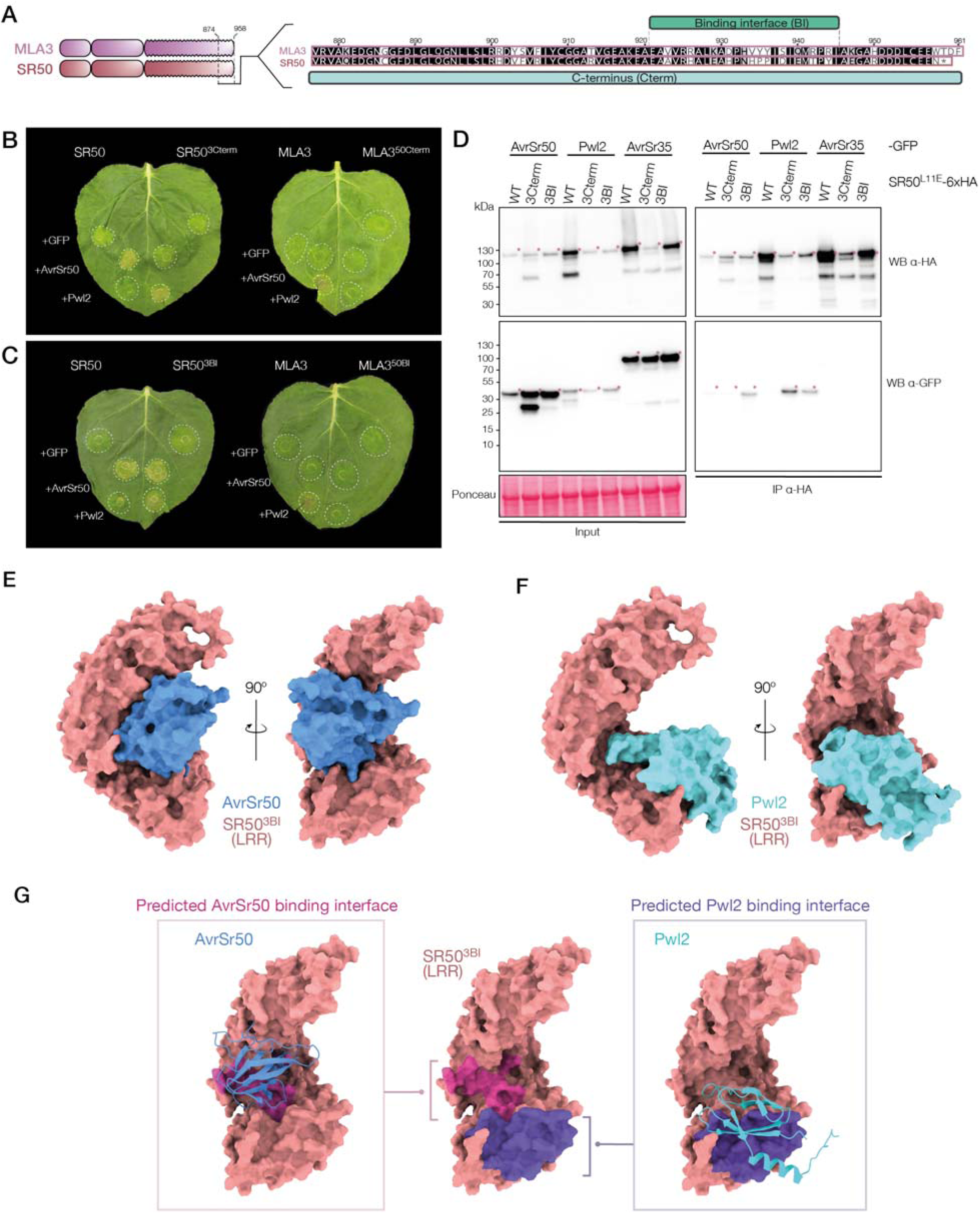
Introducing the Pwl2 binding interface into SR50 results in dual effector recognition. (**A**) Protein alignment of the C-terminus of MLA3 and SR50, schematically representing the regions swapped between MLA3 and SR50 (Cterm: C-terminus, Pwl2 binding interface: BI). (**B**) and (**C**) Cell death assays in representative leaves of *N. benthamiana* expressing the indicated proteins. Panel (B) shows MLA3-SR50 Cterm chimeras, and panel (C) shows MLA3-SR50 BI chimeras. GFP was used as negative control. The experiments consisted of three biological replicates, each with at least six technical replicates. Quantification of cell death phenotypes of (B) and (C) are shown in fig. S16. (**D**) Coimmunoprecipitation assays between SR50^L11E^ chimeric proteins and AvrSr50, Pwl2 and AvrSr35. C-terminally 6xHA-tagged SR50^L11E^ proteins were coexpressed with C-terminally 3xGFP-tagged fungal effectors. The L11E mutation in SR50 impairs cell death. AvrSr35 was used as negative control. Immunoprecipitations were done with agarose beads conjugated to HA antibodies (HA IP). Proteins were detected with the antibodies labelled on the right. Approximate molecular weights of the proteins are shown on the left. Protein loading was checked with Ponceau S solution. Asterisks show the expected protein band for each treatment. The experiment was independently repeated three times with similar results. (**E**) and (**F**) Predicted structure of the LRR domain of SR50^3BI^ in complex with AvrSr50 (E) and Pwl2 (F). The AlphaFold2 model colored by pLDDT score is shown in fig. S16. (**G**) Schematic representation of the AvrSr50 and Pwl2 binding interfaces in the LRR domain of SR50^3BI^.

## Conclusions

Our study demonstrates that MLA3 recognises the blast effector Pwl2 by mimicking the binding interface of a Pwl2 host target, HIPP43. This case of molecular mimicry represents a novel evolutionary strategy by which NLR immune receptors can acquire new pathogen recognition specificities, in addition to the known model of integration of host targets within the NLR itself as additional individual domains that serve as decoys to activate immunity (*11*) (**Fig. 5**). In contrast to the current model in which NLRs integrate effector targets to detect pathogens, our findings indicate that NLRs can achieve similar outcomes by evolving to mimic these targets. This case of host molecular mimicry by MLA3 to detect a fungal pathogen contrasts with commonly described instances of mimicry in which pathogens emulate host proteins to evade or subvert immune responses (*37*). While molecular mimicry by pathogens has been extensively studied as a strategy to manipulate host processes (*38–40*), particularly in human pathogenic bacteria (*41–44*), our findings reveal that host plant immune receptors have also evolved sophisticated mimicry strategies to counteract pathogen attack.

**Fig. 5.**
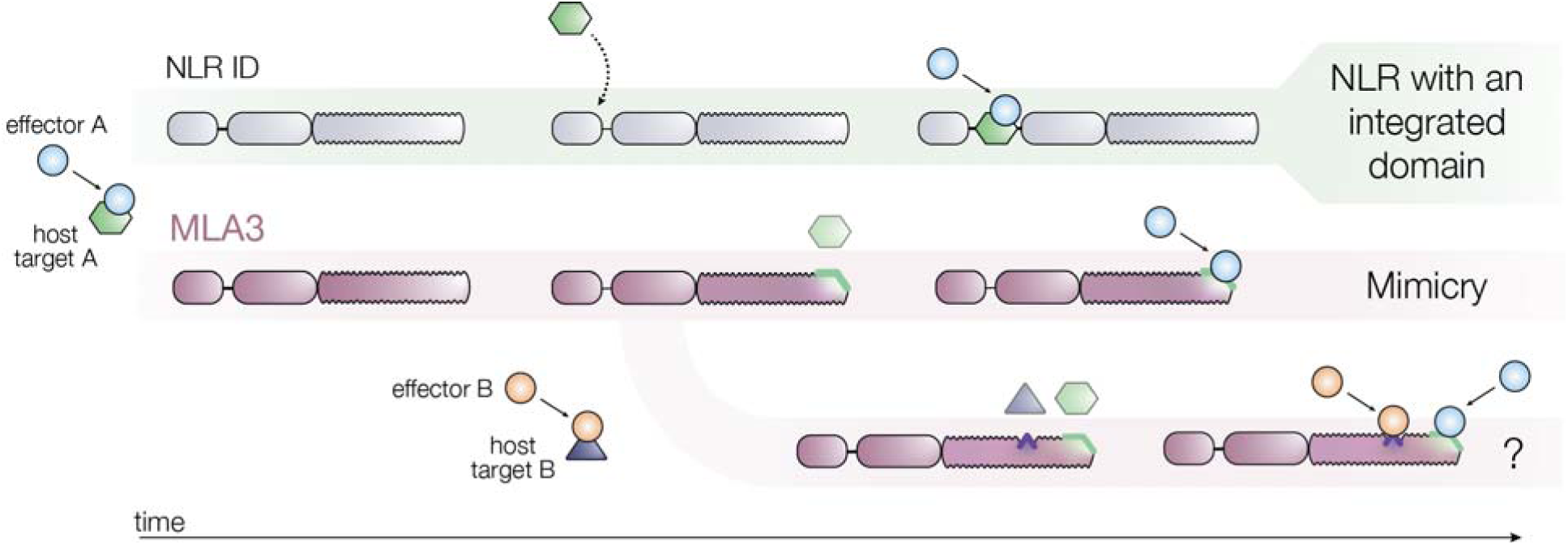
Model of evolution of molecular mimicry in MLA3 towards Pwl2 recognition. Effector targets in the host are commonly integrated into NLRs as integrated domains (IDs) that serve as baits to trigger immunity upon effector binding. We propose an alternative model for the evolution of effector recognition in which LRR domains evolve to mimic the binding interface of effector host targets to trigger immunity without integration of additional domains. This is the case for MLA3, which we propose has evolved to directly bind Pwl2 via the C-terminus of its LRR domain by mimicking the binding interface of the Pwl2 host target HIPP43. We also propose that LRR domains can contain multiple effector binding interfaces, and raise the question of whether molecular mimicry is broadly present across NLRs that directly recognize pathogen effectors and confer multiple pathogen recognition, such as MLA3.

Our findings show that the LRR domain of MLA3 binds Pwl2 through the C-terminus, diverging from the direct recognition interfaces observed in of most plant NLR resistosome structures described so far, in which effectors are recognised through the ascending lateral side of the LRR domain (*19*). Furthermore, our study also suggests that the LRR domain of NLRs can possess multiple binding interfaces, enabling them to recognize diverse pathogen effectors. This is exemplified by our bioengineering of the rye NLR, SR50. By introducing the Pwl2 binding interface of MLA3 into SR50, we generated a receptor with dual specificity: it retained its original recognition of the wheat stem rust effector AvrSr50 but also gained the ability to respond to Pwl2. This dual functionality underscores the potential of NLRs to evolve or be engineered to detect multiple pathogens through distinct binding interfaces within their LRR domains (**Fig. 5**). This might explain the ability of MLA3 to act as a resistance gene for both blast and powdery mildew. AVR_a3_, the *Bgh* effector recognised by MLA3 remains unknown. However, the *Bgh* effector repertoire is not predicted to contain effectors with a MAX fold (*45*) such as Pwl2, and most identified *Bgh* effectors recognised by other MLA alleles have an RNAse-like fold (*46*). This suggests that MLA3 may recognise structurally distinct effectors from *M. oryzae* and *Bgh*. We therefore hypothesise that the LRR domain of MLA3 might recognise Pwl2 and AVR_a3_ through distinct binding interfaces, additionally raising the question of how this would translate into different ways of receptor activation.

It remains to be addressed whether host molecular mimicry is a more common immune mechanism for pathogen detection present across the tree of life. However, our findings expand the current understanding of the molecular basis of the evolution of immune receptor specificity, and pave the way for further exploration into the bioengineering and targeted modification of NLRs to enhance crop resistance against multiple pathogens.

## Acknowledgments

We thank J. Rhodes for his insightful questions and suggestions, which greatly shaped the direction of this story. We are very thankful to H.J. Brabham and V. Were, who laid the groundwork for this project. We also thank S. Kamoun, Y. Sugihara, A. Toghani, A. Posbeyikian and D. Lüdke for the useful discussions. D.G.D.L.C. thanks M. Bergum, P. Green and I. Hernández-Pinzón for support and encouragement. We thank all members of TSL support teams for their invaluable assistance.

## Funding

The Gatsby Charitable Foundation, supporting MJM and NJT

UKRI-BBSRC grant APH-ISP BB/X010996/1, supporting MJB, MJM and NJT.

UKRI-BBSRC grant BB/W00108X/1, supporting RZ and MJB.

United States Department of Agriculture-Agricultural Research Service CRIS #5062-21220-025-000D, supporting MJM

## Author contributions

Conceptualization: DGDLC, MJM, NJT.

Methodology: DGDLC, MJM, NJT.

Investigation: DGDLC, RZ.

Visualization: DGDLC, MJM.

Funding acquisition: MJM, NJT, MJB.

Project administration: MJM, NJT.

Supervision: MJM, NJT

Writing – original draft: DGDLC, MJM, NJT.

Writing – review & editing: DGDLC, MJM, NJT, RZ, MJB.

## Data and materials availability

Sequencing data used in this study are found in the NCBI database under BioProject codes PRJNA761551, PRJNA378723, and PRJNA1138684. All other data are available in the main text or the supplementary materials. A material transfer agreement with The Sainsbury Laboratory is required to receive plasmids. The use of the materials will be limited to noncommercial research uses only. Please contact N.J.T. regarding biological materials, and requests will be responded to within 60 days.

## Materials and Methods

### Plant material and growth conditions

Barely (*Hordeum vulgare*) accessions for infection assays were grown in a Sanyo cabinet at 25°C and a 16-h photoperiod. Wild-type *Nicotiana benthamiana* plants were grown in a controlled environment room at 22°C, humidity range of 45% to 65% and a 16-h light cycle.

### *Magnaporthe oryzae* growth and inoculation

*M. oryzae* wild-type isolate KEN54-20 and derived *pwl2* mutants (*15*) were routinely grown on complete medium agar plates at 24 C with a 12-h light cycle. For long term storage, isolates were grown over sterile Whatman filter paper (GE Healthcare Whatman™ Qualitative Filter Paper, Fisher Scientific UK) placed on top of CM agar plates. Filter papers were subsequently dehydrated and stored at -20°C. For infection assays, conidia were collected from one week old *M. oryzae* culture plates by adding 5 mL of sterile dH_2_O and scraping the mycelia with the tip of a 1.5 mL microcentrifuge tube. The conidial suspension was filtered through Miracloth^TM^ and collected in a 50mL collection tube. The supernatant was discarded, and the conidia pellet was resuspended in 5 mL of 0.2% (w/v) gelatin. Conidia concentration was determined using a haemocytometer and adjusted to 1x10^5^ conidia mL^-1^. For leaf drop inoculations on barley, the first leaf of 7-day old seedlings was detached and placed on agar boxes (5 g/L agar-agar, 0.1 g/L benzimidazole). Each leaf was inoculated with 3 to 4 drops of 5 µL of the conidial suspension. Agar boxes with inoculated barley leaves were placed in a growth cabinet at 25°C and a 16/8 h light/dark cycle. Infection phenotypes were recorded 7 days post inoculation (dpi).

### Transcriptome analysis of *Mla* in barley accessions Baronesse and S13

Paired end Illumina RNA-seq was trimmed using Trimmomatic (v0.36) using parameters: ILLUMINACLIP:TruSeq3-PE.fa:2:30:10 LEADING:5 TRAILING:5 SLIDINGWINDOW:4:10 MINLEN:36. Alignment of Illumina RNA-seq from barley accessions Baronesse (PRJNA378723/SRR5345035) and S13 (PRJNA761551/SRR15828851) to the *Mla3* genomic region (*15*) was performed using hisat2 (v2.2.1) using default parameters. Data visualization was performed in Geneious Prime (v2023.0.4). First leaf tissue was harvested from barley accessions Baronesse and S13 at 10 d after sowing grown in the greenhouse under natural daylight conditions. Tissue was flash frozen in liquid nitrogen, stored at −80 °C, and homogenized into a fine powder in liquid nitrogen–chilled pestle and mortars. RNA extraction was performed using TRI-reagent (Sigma-Aldrich; T9424) according to the manufacturer’s protocol. Samples were treated with RQ1 RNase free DNase (Promega; M6101) to remove residual DNA, purified using RNeasy mini spin columns (Qiagen; product No. 74104), and assessed using RNA Nano Chips (Agilent Technologies; product no. 5067-1511) on an Agilent 2100 Bioanalyzer. Sequencing libraries for Oxford Nanopore Technology sequencing were prepared using the cDNA-PCR kit (Oxford Nanopore Technology; SQK-PCS109) according to the manufacturer’s protocol. Libraries were applied to individual MinION flow cells (Oxford Nanopore Technology; R9.4.1). Raw reads were trimmed using Porechop (v0.2.4) with default parameters. *De novo* transcriptome assembly was performed using RNA-Bloom (v2.0.1) with default parameters. BLAST (v2.12.0+) onto assemblies was performed using *Mla3* and *Mla23* as query using default parameters.

### Plasmid constructions

The constructs of C-terminally HA-tagged MLA3, MLA3^L11E^, and FLAG-tagged Pwl2 used in this study are the same as those used in our previous report (*15*). The coding sequences of *Mla23* (ACZ65493.1) and *Sr50* (QNU41030.1) were domesticated to remove internal *BsaI* and *BpiI* restriction sites, codon optimised for expression in *N. benthamiana*, and synthesized by Twist Bioscience (San Francisco, CA, United States). The synthesised *Mla23* and *Sr50* fragments did not contain a stop codon, and were cloned in the pTwist_Kan_High_Copy cloning vector (Twist Bioscience) with flanking *BsaI* restriction sites for subsequent Golden Gate assemblies (*47*). New *Mla3*, *Mla23* and *Sr50* constructs, either carrying specific mutations or containing smaller fragments, were obtained through polymerase chain reaction (PCR) amplification using Phusion^TM^ High-fidelity DNA polymerase (ThermoFisher Scientific), according to the manufacturer’s instructions. Primers to PCR amplify *Mla3* fragments are listed in **Table S2**. PCR fragments were purified and cloned into the level 0 acceptor plasmid pICSL01005 (TSL SynBio) using Golden Gate cloning.

For transient gene expression assays in *N. benthamiana,* coding sequences were cloned into the level one binary acceptor pICH47732 (Addgene no. 4800) by Golden Gate assemblies. *Mla3* and *Mla23* variants were cloned with pICH85281 (mannopine synthase + Ω promoter (Mas Ω), Addgene no. 50272), pICSL50009 (6xHA C-terminal tag, Addgene no. 50309), and pICSL60008 (Arabidopsis heat shock protein terminator (AtHSP18 terminator), TSL SynBio). *Mla3* and *Mla23* mutant versions were generated by inverse PCR using the primers listed in **Table S3** and **Table S4**, and the corresponding *Mla* wild-type allele of interest as template. The PCR products were digested with *BsaI* and religated. Substitution of residue K926 in MLA3 for all possible 19 amino acids was done by annealing pairs of oligos listed in **Table S5** containing the corresponding K926 mutation with flanking *BsaI* restriction sites. Briefly, matching oligos were mixed in equimolar concentrations and incubated at 95°C for 2 min and gradually cooled down to 25°C over 45 min. In parallel, the *Mla3* sequence outside the region covered by the oligos and embedded in the level one assembly for transient expression was amplified by inverse PCR using the primers listed in **Table S5**. The purified PCR product and each pair of annealed oligos was assembled by Golden Gate cloning using *BsaI. Sr50-Mla3* chimeras were synthesised by GENEWIZ standard gene synthesis (Azenta Life Sciences). *Sr50* and *Mla3* fragments were cloned into the binary acceptor pICH47732, with pICH87644 (AtAct2 + Ω promoter, Addgene no. 50274), pICSL50009 (6xHA C-terminal tag), and pICSL60008 (AtHSP18 terminator), TSL SynBio).

The coding sequence of AVR-PikD was PCR amplified from a previously cloned construct (*48*) and cloned into the level 0 acceptor plasmid pICSL01005 (TSL SynBio) using Golden Gate cloning. For transient expression assays in *N. benthamiana,* AVR-PikD was cloned into the binary acceptor plasmid pICH47732 with pICH51266 (long 35S + Ω promoter, Addgene no. 50267), pICSL50007 (3xFLAG C-terminal tag, Addgene no. 50308), and pICH41414 (35S terminator, Addgene no. 50337). The coding sequences of *AvrSr50* and *AvrSr35* were codon optimised for expression in *N. benthamiana,* synthesised by GENEWIZ standard genes synthesis (Azenta Life Sciences) without internal *BsaI* or *BpiI* restriction sites, and *BsaI* flanking sites for Golden Gate assemblies. The level 0 plasmid containing the coding sequence of *PWL2* was cloned previously (*15*). *PWL2, AvrSr50* and *AvrSr35* were cloned into the binary acceptor plasmid pICH47732 with pICH51266 (long 35S + Ω promoter), pICSL50008 (GFP C-terminal tag, Addgene no. 50314), and pICH41414 (35S terminator).

The coding sequences of AtTRXh5 (Q39241.1) and OsHIPP43-HMA (LOC_Os01g32330.1) were synthesised as gene fragments by Twist Bioscience (San Francisco, CA, USA) domestication to remove internal *BsaI* and *BpiI* restriction sites. Flanking adapters with *BpiI* restriction sites were added for cloning into the pICSL01005 (TSL SynBio) acceptor plasmid using Golden Gate cloning. For transient expression in *N. benthamiana,* the AtTRXh5 and OsHIPP43-HMA level 0 modules were cloned into the level 1 binary acceptor plasmid pICH47732 with pICSL13001 (long 35s CaMV promoter + 5’ untranslated leader, Addgene no. 50265), pICSL30009 (4xMyc N-terminal tag, Addgene no. 50301), and pICH41414 (35S terminator). The module pICSL50028 (TSL SynBio) was included to add a stop codon.

For yeast-two-hybrid assays, all the relevant coding sequences cloned in the level 0 acceptor pICSL01005 were cloned into the level one acceptor plasmids pGADT7 or pGBKT7 (TSL SynBio) by Golden Gate assembly. The module pICSL50028 (TSL SynBio) was included to add a stop codon.

### *Agrobacterium-*mediated transient protein expression in *N. benthamiana*

Transient gene expression in *N. benthamiana* for both cell death and coimmunoprecipitation assays was performed by delivering T-DNA constructs transformed in *A. tumefaciens* GV3101::pMP90 into the leaves of 4-week-old *N. benthamiana* plants. Overnight cultures of *A. tumefaciens* carrying the constructs of interest were grown in LB medium at 28°C with constant shaking. After centrifugation at 5000xg for 10 minutes, cell pellets were resuspended in infiltration buffer (10 mM MES, 10 mM MgCl2, and 150 µM acetosyringone). The OD_600_ of each bacterial suspension was measured and adjusted to a final working concentration of 0.5 for *Mla* constructs, 0.25 for constructs encoding *Sr50* and 0.4 for constructs encoding fungal effectors. The third and fourth leaves from the top of the 4-week-old *N. benthamiana* plants were infiltrated using a 1 mL needleless syringe. For coimmunoprecipitation assays, two entire leaves were infiltrated per treatment and harvested 3 days post-infiltration. For cell death assays, the appropriate combinations of *A. tumefaciens* were spot infiltrated as specified in each experiment, and phenotypes were recorded 5-6 days post-infiltration.

### Protein extraction and coimmunoprecipitation assays

Two *N. benthamiana* leaves were collected 3 days post-agroinfiltration, flash-frozen in liquid nitrogen, and ground using a Geno/Grinder tissue homogenizer. The ground leaf tissue was homogenized with GTEN extraction buffer (10% glycerol, 25 mM Tris-HCl pH 7.5, 1 mM EDTA, and 300 mM NaCl) at a 1:2 weight/volume ratio, supplemented with 2% PVPP, 10 mM DTT, 1% protease inhibitor cocktail (Sigma), and 0.2% IGEPAL CA-630. The samples were centrifuged at 5,000 × g for 20 minutes at 4°C, and the supernatant was collected and centrifuged again under the same conditions for an additional 10 minutes. The final supernatant was filtered through a 0.45-µm Minisart filter to obtain the total protein extract.

For coimmunoprecipitation assays, 1 mL of the total protein extract was incubated with 20 μL of Anti-HA Affinity Matrix (rat IgG1, Roche) or Anti-FLAG M2 Affinity Gel (Sigma-Aldrich) by end-over-end mixing for 90 minutes at 4°C. The beads were then washed five times with immunoprecipitation washing buffer (GTEN extraction buffer with 0.2% IGEPAL CA-630) and resuspended in 60 μL of 1× Laemmli sample buffer (BioRad) with 10 mM DTT. Proteins were eluted from the beads by heating the samples at 80°C for 10 minutes.

Total protein extracts and immunoprecipitated samples were separated by SDS-PAGE on 4-20% BioRad Mini-PROTEAN TGX gels, alongside a PageRuler Plus prestained protein ladder (Thermo Scientific), and transferred to a PVDF membrane using the Trans-Blot Turbo transfer system (BioRad) according to the manufacturer’s instructions.

### Immunoblotting

PVDF membranes were blocked for 60 minutes in a solution of 5% skim milk powder in 1× TBST. Epitope tagged proteins were detected with Anti-HA-Peroxidase, high-affinity antibody from rat IgG1 (12013819001; Roche), ANTI-FLAG M2-Peroxidase (HRP) antibody produced in mouse (A8592; Sigma-Aldrich), or Anti-GFP Antibody (B-2) HRP (sc-9996 HRP; Santa Cruz), each diluted 1:5,000 in 5% skim milk powder in 1× TBST. Proteins were detected by adding a 1:1 mix of Pierce ECL Western Blotting Substrate and SuperSignal West Femto Maximum Sensitivity Substrate (Thermo Fisher Scientific). Membranes were imaged using an Amersham ImageQuant 800 western blot imager system. Protein loading was verified by staining with Ponceau S solution (Sigma).

### Cell death assays

Cell death assays to test hypersensitive response (HR) were carried out by transiently expressing the genes of interest in leaves of 4 week old *N. benthamiana* plants as describe above. The cell death phenotypes were scored 3 dpi using the previously published cell death phenotypic scale (Maqbool, 2015) ranging from 0: no response, to 7: confluent cell death. Statistical analysis of the quantified cell death phenotypes was performed by implementing the besthr R library (*49*)

### Yeast-two-hybrid assays

Chemically competent *Saccharomyces cerevisiae* Y2HGold cells (Takara Bio) were co transformed with the pGBKT7 and pGADT7 constructs containing the genes of interest, using the commercial kit Frozen-EZ Yeast Transformation II^TM^ (Zymo Research) following the manufacturer’s instructions. Co-transformed yeast cells were selected on SD solid medium not supplemented with leucine or tryptophane (SD-LW) after incubation at 28°C for three days. Single yeast colonies grown on SD-LW plates were used to inoculate liquid SD-LW medium and incubated overnight at 28°C and 190 rpm. Saturated cultures were adjusted to an OD_600_ of 1 and used for three tenfold serial dilutions (10^-1^, 10^-2^ and 10^-3^). 5 µL drops of the initial yeast suspensions and each dilution were placed on SD-LW agar plate, and on SD solid media not supplemented with leucine, tryptophane, histidine or adenine (SD-LWHA) containing X-a-gal. Yeast growth was documented after incubation at 28°C for four days.

### Protein complex modelling

Protein structural predictions were performed using Alphafold2 (v2.3.1) with a PDB selection date threshold of 2023-07-26 and monomer or multimer models when input included one or two proteins, respectively. Visualization of protein structures was performed using ChimeraX (1.8rc202405310011). Protein structural alignments were performed in ChimeraX using the Matchmaker tool using default parameters.

### Phylogenetic analyses

Multiple sequence alignment of the *RGH1* gene family was performed using MUSCLE (v5.1) with codon-based approach. Phylogenetic analysis was performed using RAxML (v8.2.12) using the general time reversible model of nucleotide substitution under the gamma model of rate heterogeneity. *Brachypodium distachyon Mla* orthologs were used as outgroups (*Bradi2g39247.4* and *Bradi2g39393.1*). Bootstrap support was estimated based on 1,000 bootstraps. Tree visualization was performed using iTOL (https://itol.embl.de/).

---

**Fig. S1.**
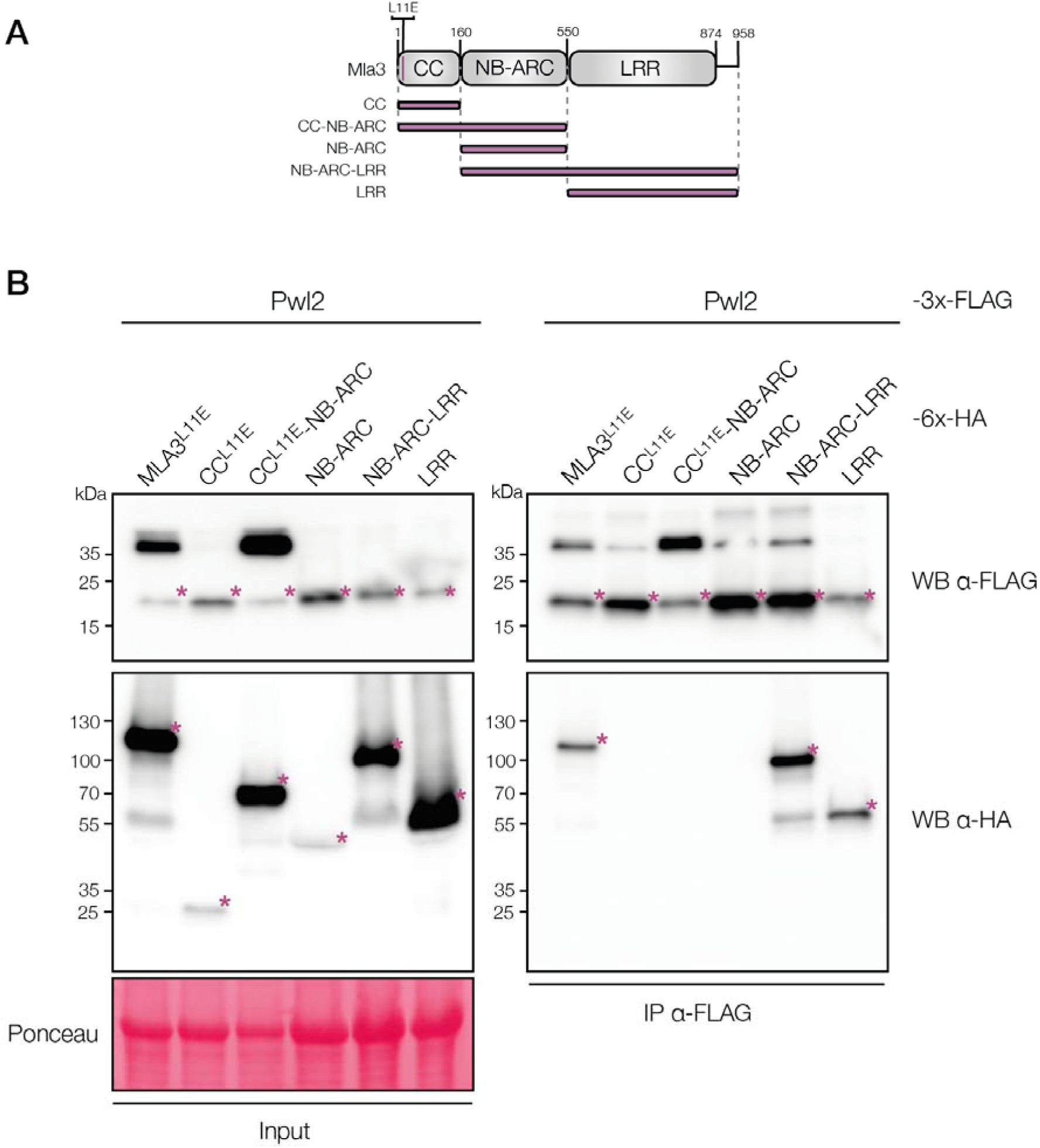
The LRR domain of MLA3 is required and sufficient for association with Pwl2 in *N. benthamiana*. (A) Schematic diagram of the domain architecture of MLA3 and the boundaries used to delimit the truncations for the constructs used in co-immunoprecipitation assays. (B) Coimmunoprecipitation assay done by coexpressing 6xHA C-terminally tagged Mla3^L11E^ (115.6 kDa), CC^L11E^ (25.7 kDa), CC^L11E^-NB-ARC (70 kDa), NB-ARC (52 kDa), NB-ARC-LRR (97.5 kDa) or LRR (53.2 kDa) with 3xFLAG C-terminally tagged Pwl2 (17.3 kDa) in *N. benthamiana*. Immunoprecipitation (IP) was performed with anti-FLAG agarose beads. Total protein (Input) and pulled-down fractions (IP) were detected with the antibodies labelled on the right. Asterisks show the expected protein band for each treatment. Protein loading was checked with Ponceau S solution (Sigma). The experiment was independently repeated three times with similar results.

**Fig. S2.**
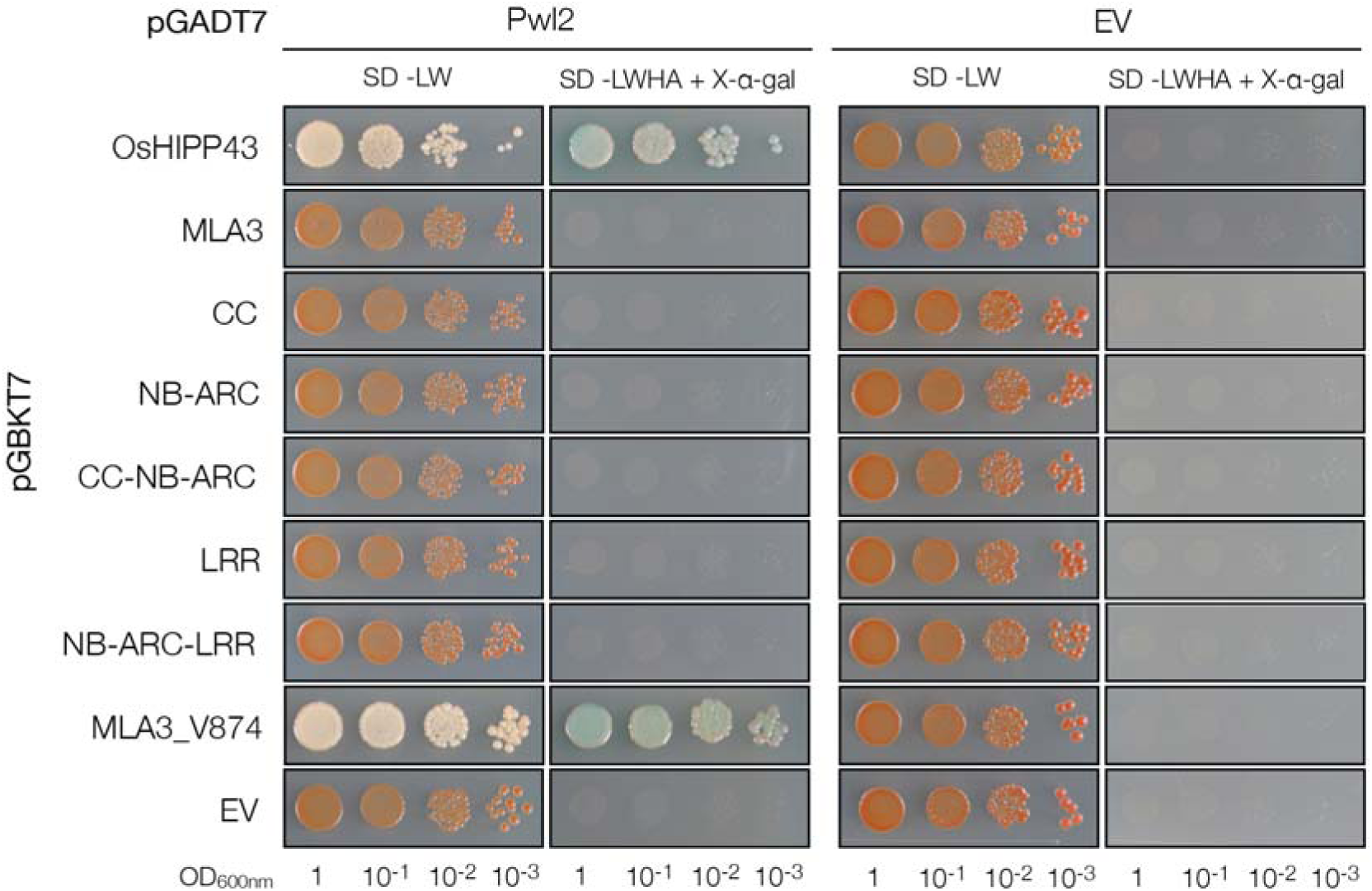
The last 85 residues of MLA3 are sufficient to interact with Pwl2 in yeast. Yeast-two-hybrid assay to test association between MLA3 fragments (CC, CC-NB-ARC, NB-ARC, NB-ARC-LRR, LRR and MLA3_V874) and Pwl2. Yeast cells were co-transformed with the indicated constructs. The plasmids pGBKT7 and pGADT7 contain the Gal4 binding domain and the Gal4 activating domain, respectively, fused to the indicated gene. OsHIPP43 was used as positive control for interaction with Pwl2. Yeast growth on SD–LW media indicates co-transformation with both indicated plasmids. Yeast growth on SD–LWHA + X-gal media indicates positive protein interactions. Photos were taken after 4 days of growth at 28 . The experiment was repeated two times with similar results.

**Fig. S3.**
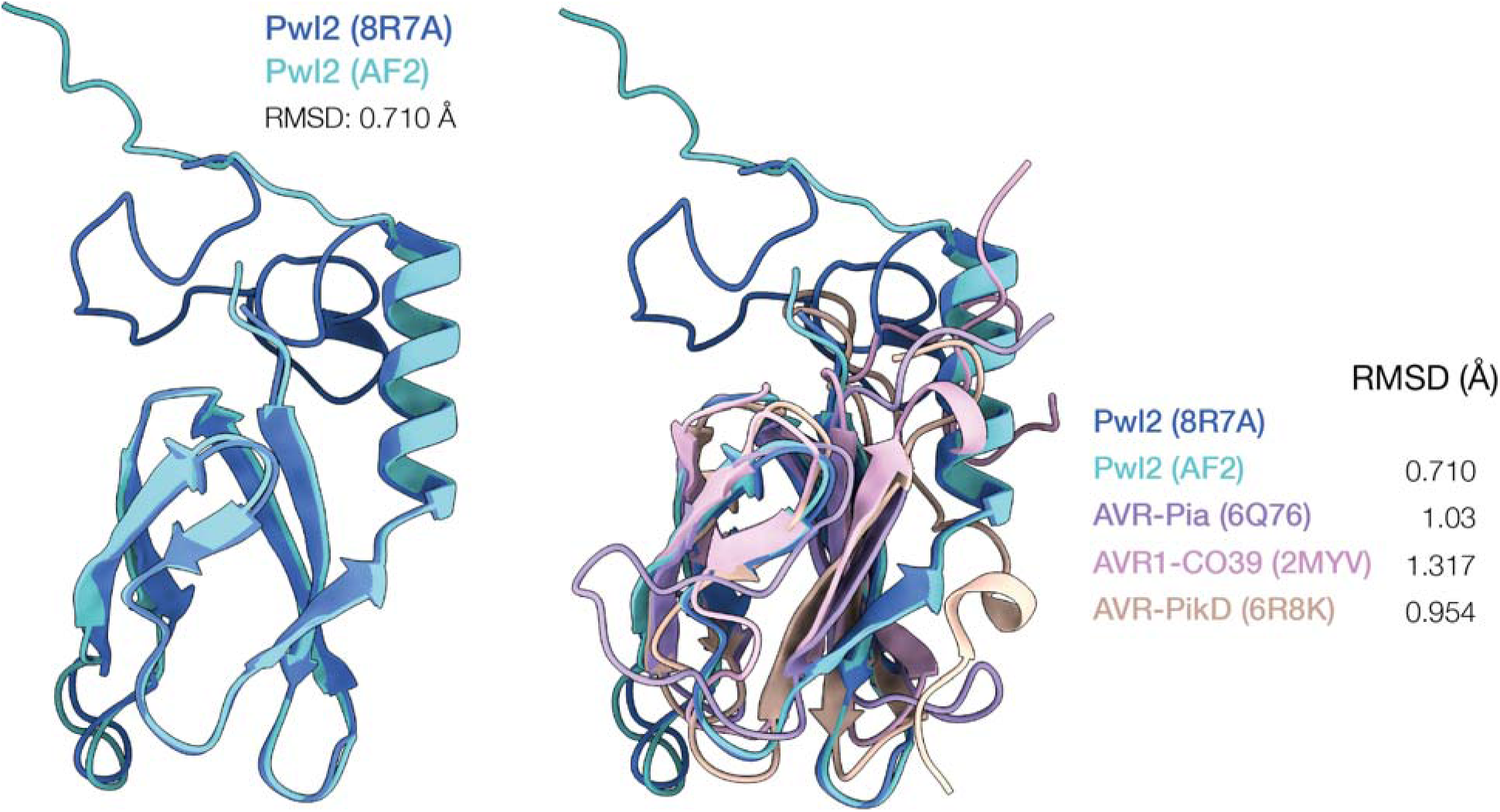
The structure of Pwl2 is accurately predicted by AlphaFold2. Superimposition of the X-ray crystal structure of Pwl2 (PDB ID: 8R7A) with the AlphaFold2 (AF2) structure prediction of Pwl2 (left). AF2 correctly predicts that Pwl2 is a MAX effector. Known MAX effectors (AVR-Pia (PDB ID: 6Q76), AVR1-CO39 (PDB ID: 2MYV), and AVR-PikD (PDB ID: 6R8K)) are superimposed with the Pwl2 AF2 structure and compared to the X-ray crystal structure of Pwl2 (right).

**Fig. S4.**
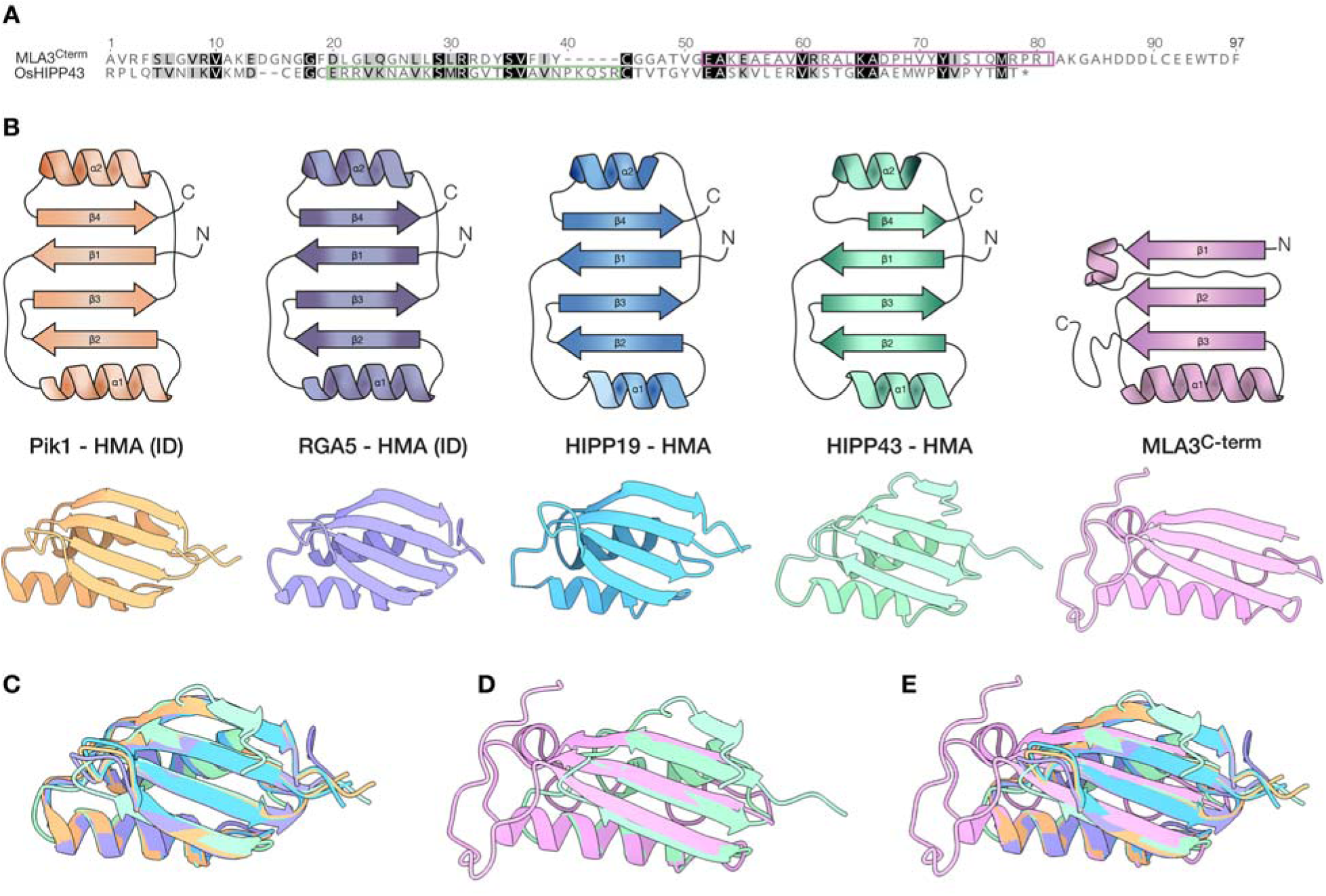
The C-terminus of MLA3 does not have the structure of a Heavy Metal Associated (HMA) domain. (**A**) Protein sequence alignment of the C-terminus of MLA3 and HIPP43. The Pwl2 binding interface is marked with a pink or green box in MLA3 and HIPP43, respectively. (**B**) Secondary and tertiary structure of HMA domains (Pik1 HMA - PDB: 6R8K; RGA5 HMA - PDB: 5ZNE; HIPP19 HMA - PDB: 7B1I, HIPP43 HMA - PDB: 87RA) and the C-terminus of MLA3, predicted by AlphaFold2 (AF2). ID: Integrated domain. (**C**) Superimposition of the structure of the HMA domains shown in (B). (**D**) Superimposition of the HMA domain of HIPP43 with AF2 predicted structure of the C-terminus of MLA3. (**E**) Superimposition of the protein structured shown in (B).

**Fig. S5.**
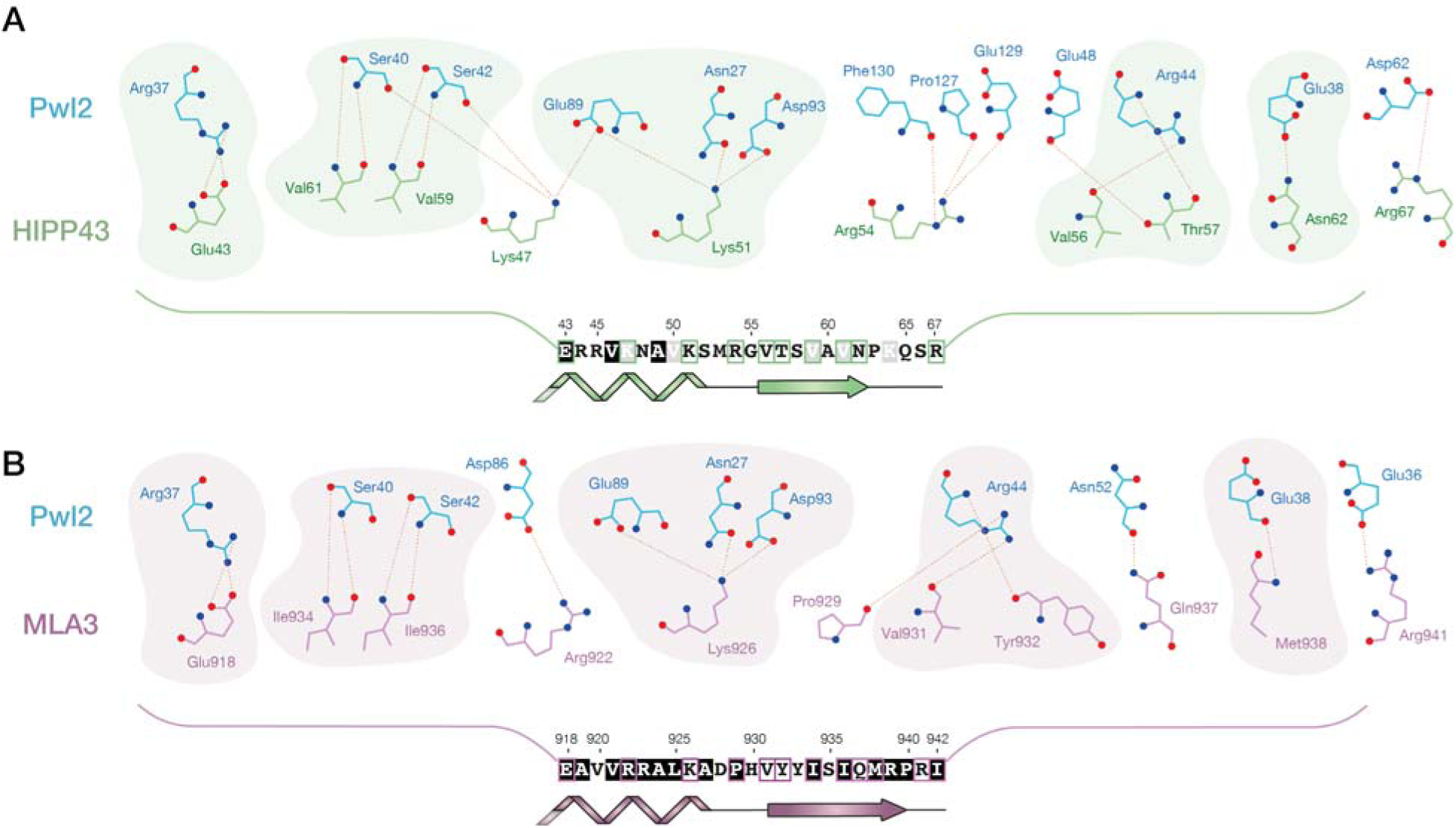
Schematic comparison of the Pwl2 binding interface in OsHIPP43 and MLA3. (**A**) Amino acids of OsHIPP43 (green) that interact with Pwl2 (blue). Yellow dotted lines show non-covalent bonds between Pwl2 and HIPP43. The amino acid sequence of HIPP43 at the binding interface is shown. Residues that interact with Pwl2 are highlighted in green boxes. Green shading of interacting amino acids denotes similarity or conservation with MLA3. (**B**) Amino acids of MLA3 (pink) that interact with Pwl2 (blue). Yellow dotted lines show non-covalent bonds between Pwl2 and MLA3. The amino acid sequence of MLA3 at the binding interface is shown. Residues that interact with Pwl2 are highlighted in pink boxes. Pink shading of interacting amino acids denotes similarity or conservation with HIPP43.

**Fig. S6.**
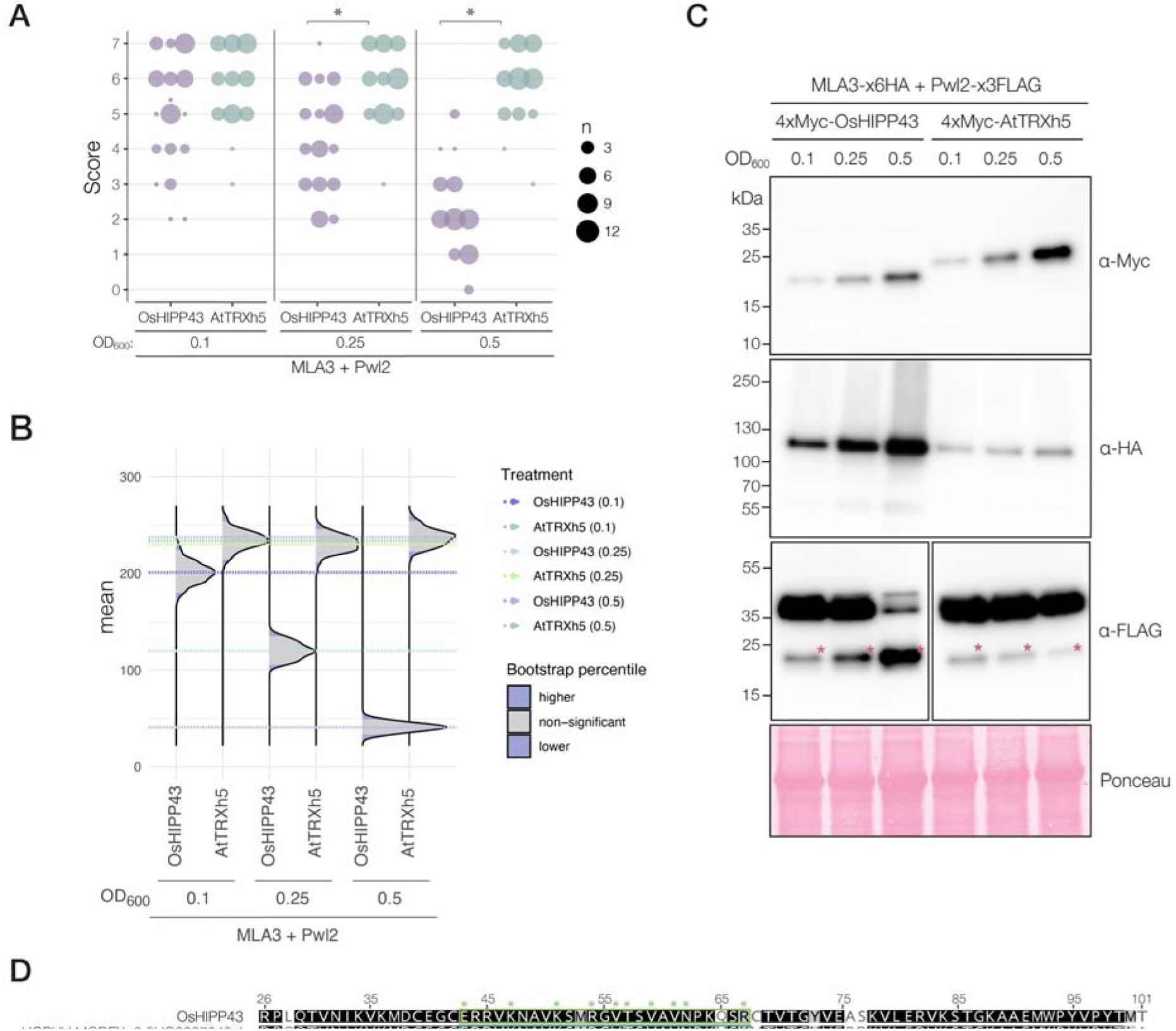
Quantification of cell death phenotypes of Fig. 2D. Cell death phenotypes from Fig. 2D were scored at three days post-agroinfiltration. The results are presented as a dot plot, where the size of a dot is proportional to the number of technical samples with the same score within the same biological replicate. The experiments were independently repeated three times with at least 6 technical replicates; the three data point columns of each tested condition correspond to results from different biological replicates. Significant differences between conditions were calculated based on bootstrapping rank statistics and are indicated with an asterisk. (**B**) Statistical analysis of the cell death response shown in (A) performed with the estimation method from the besthr R library (*49*). Distribution of 1000 bootstrap sample rank means, with the areas in blue highlighting the 0.025 and 0.975 percentiles of the distribution. The grey area indicates the 95% confidence interval of the ranked mean. The difference of cell death response is considered statistically significant if the confidence interval of the ranked mean of a given treatment falls outside of the confidence interval of another condition (i.e. within or beyond the blue percentile of the mean distribution of another condition). (**C**) Western blots to confirm protein expression in Fig. 2D using the antibodies labelled on the right. MLA3 was C-terminally tagged with 6xHA (115.6 kDa), Pwl2 was C-terminally tagged with 3xFLAG (17.3 kDa), and OsHIPP43 and AtTRXh5 were N-terminally tagged with 4xmyc (16 kDa, and 18 kDa, respectively). Total protein extracts were prepared from leaves harvested 3 days after agroinfiltration. Protein loading was checked with Ponceau S solution (Sigma) Asterisks show the expected size of the protein of interest. (**D**) Protein sequence alignment of the HMA domain in OsHIPP43 and the HMA domain of the closest homologue in barley, HORV.MOREX.r3.3HG0267040.1. The alignment is colored based on amino acid similarity. The Pwl2 binding interface is highlighted inside the green box. Residues of OsHIPP43 that interact with Pwl2 are marked with an asterisk.

**Fig. S7.**
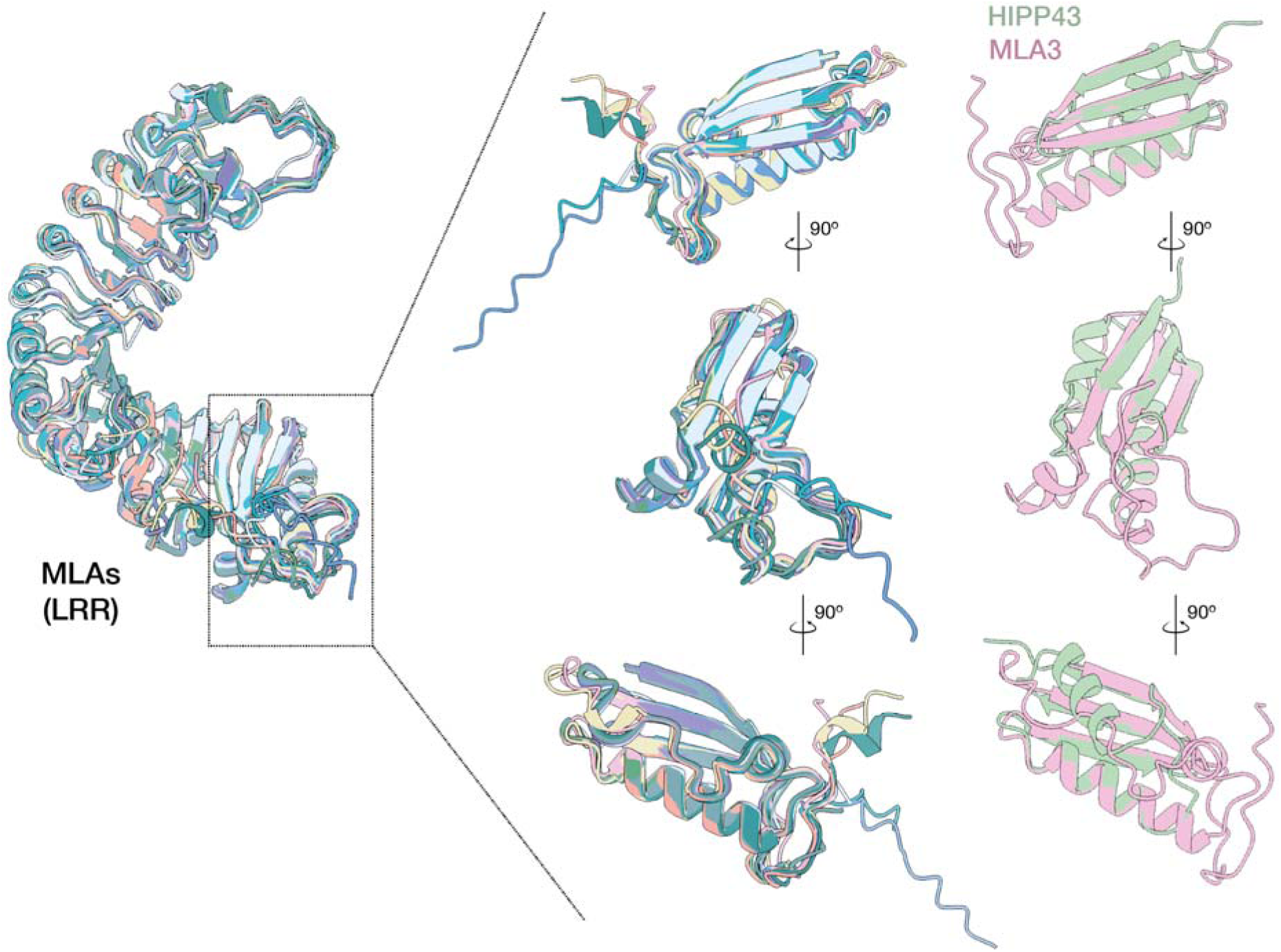
The structure of the LRR domain of MLA alleles is conserved. Superimposition of the AlphaFold2 structure prediction of the LRR domain of representative MLA alleles from fig. 3A and MLA3 (pink). The C-terminus of the MLA alleles is zoomed in, and compared to the C-terminus of MLA3 (pink) and the structure of HIPP43 (green).

**Fig. S8.**
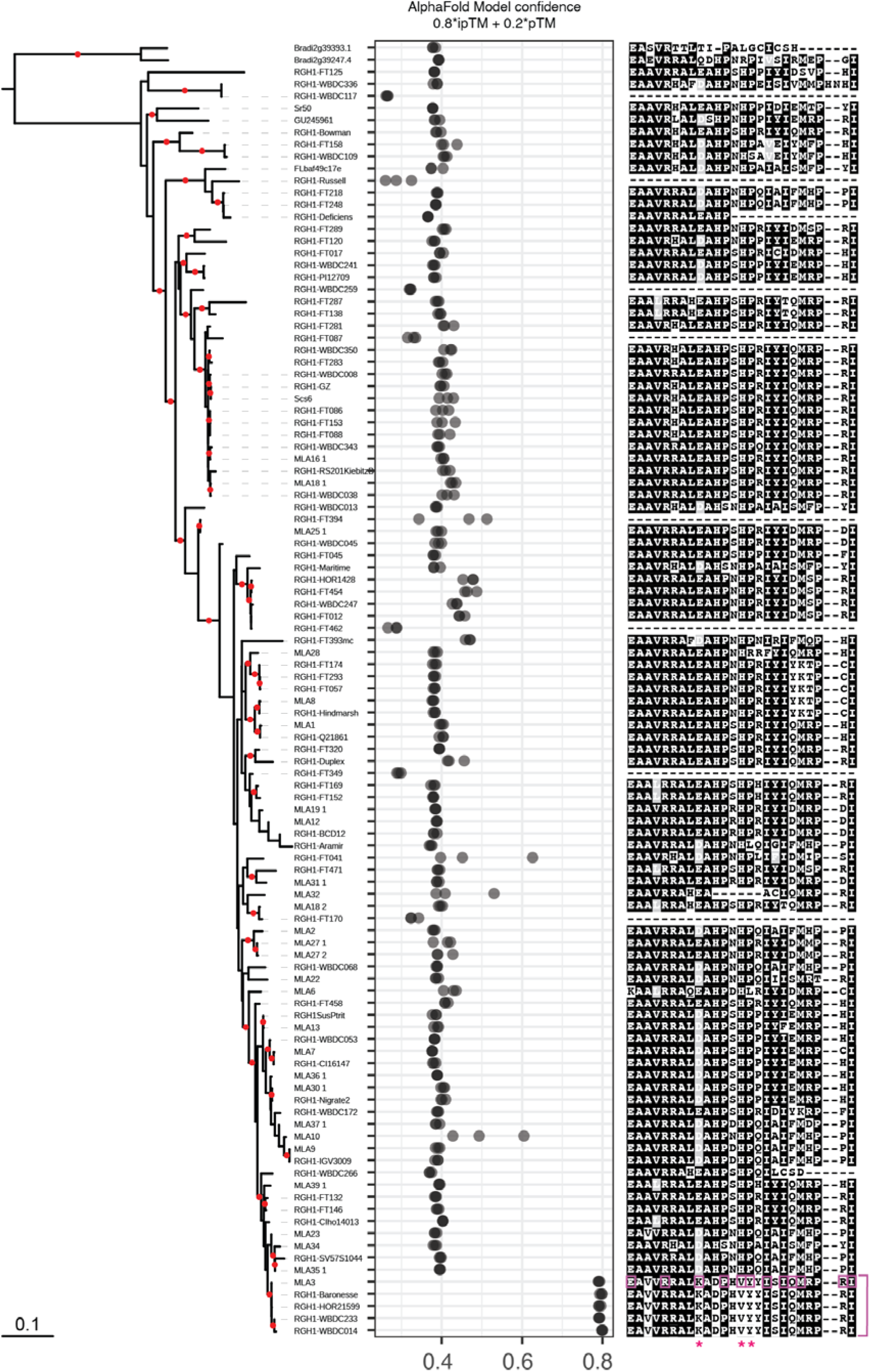
MLA3 and MLA3-like alleles are uniquely predicted to interact with Pwl2. Phylogenetic tree based on a codon-based alignment of the full sequence of all MLA alleles. Branches with bootstrap support >80 are marked with a red dot. The corresponding AlphaFold model confidence score (0.8*ipTM + 0.2*pTM) of the LRR domain each MLA allele in complex with Pwl2 is plotted in front of the tree. Structure predictions were performed three independent times. Each dot represents the confidence score of each round of prediction. The multiple protein sequence alignment of the region corresponding to the Pwl2 binding interface in MLA3 is shown. MLA3 residues that interact with Pwl2 are highlighted in pink boxes. Amino acids that are unique in MLA3 at the Pwl2 binding interface are shown with an asterisk (K926, V931 and Y932). The clade of MLA3-like alleles, with a C-terminus identical to MLA3 is highlighted in pink, at the bottom of the tree.

**Fig. S9.**
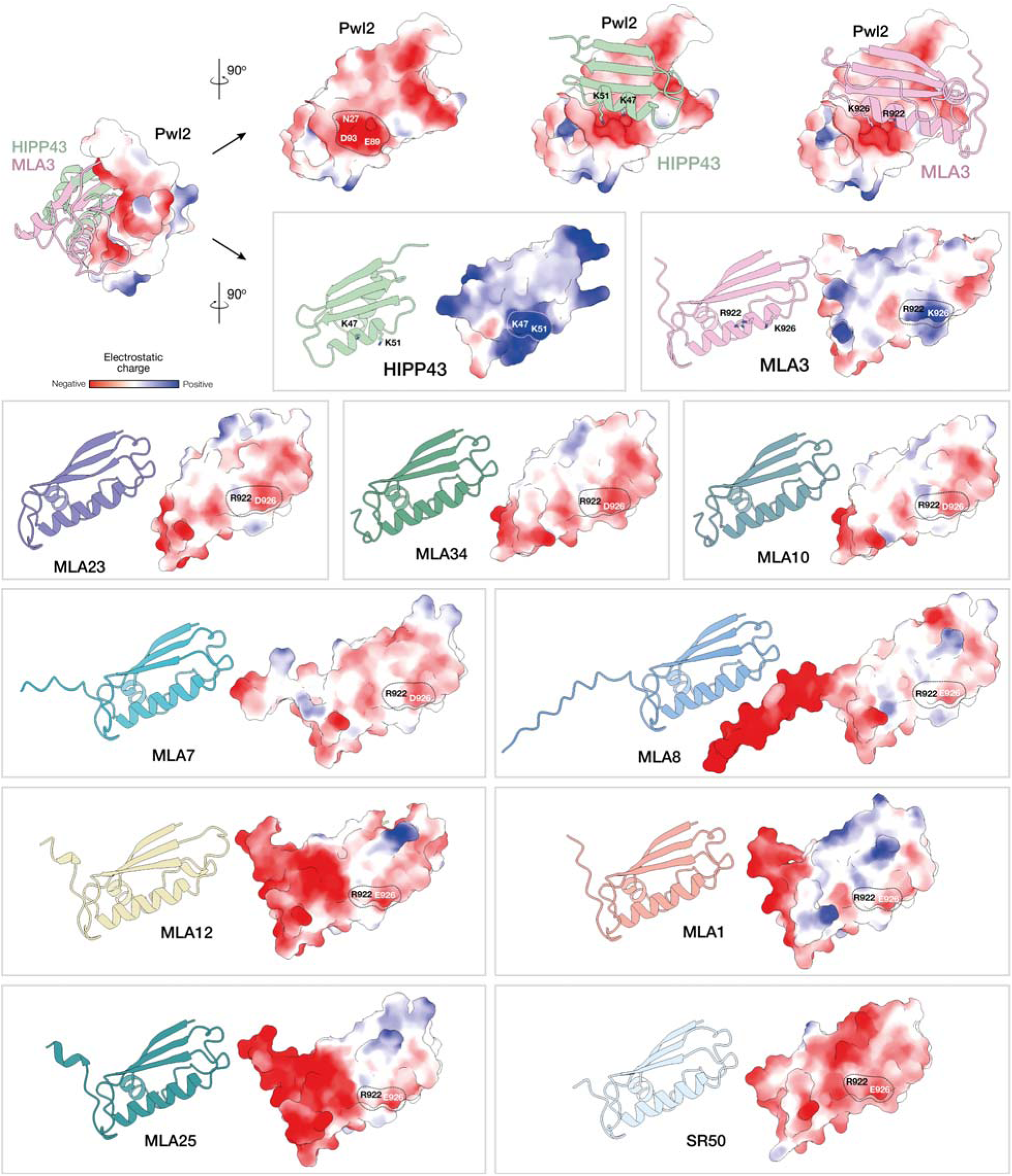
MLA3 has a unique positive charge in the Pwl2 binding interface that mimics HIPP43. Superimposition of the predicted structure of Pwl2 in complex with the C-terminus of MLA3 and the structure of the Pwl2–HIPP43 complex (Top left corner). The surfaces of Pwl2, MLA3 and HIPP43 are colored according to the electrostatic charge at the binding interface. The matching residues K47 and K51 in HIPP43, and R922 and K926 in MLA3 create a positively charged patch that interacts with a negative pocket in Pwl2 made by the amino acids N27, E89 and D93. The presence of an acidic residue (E or D) instead of K926 in all other MLA alleles abolishes the interaction with Pwl2 due to lack of charge complementarity.

**Fig. S10.**
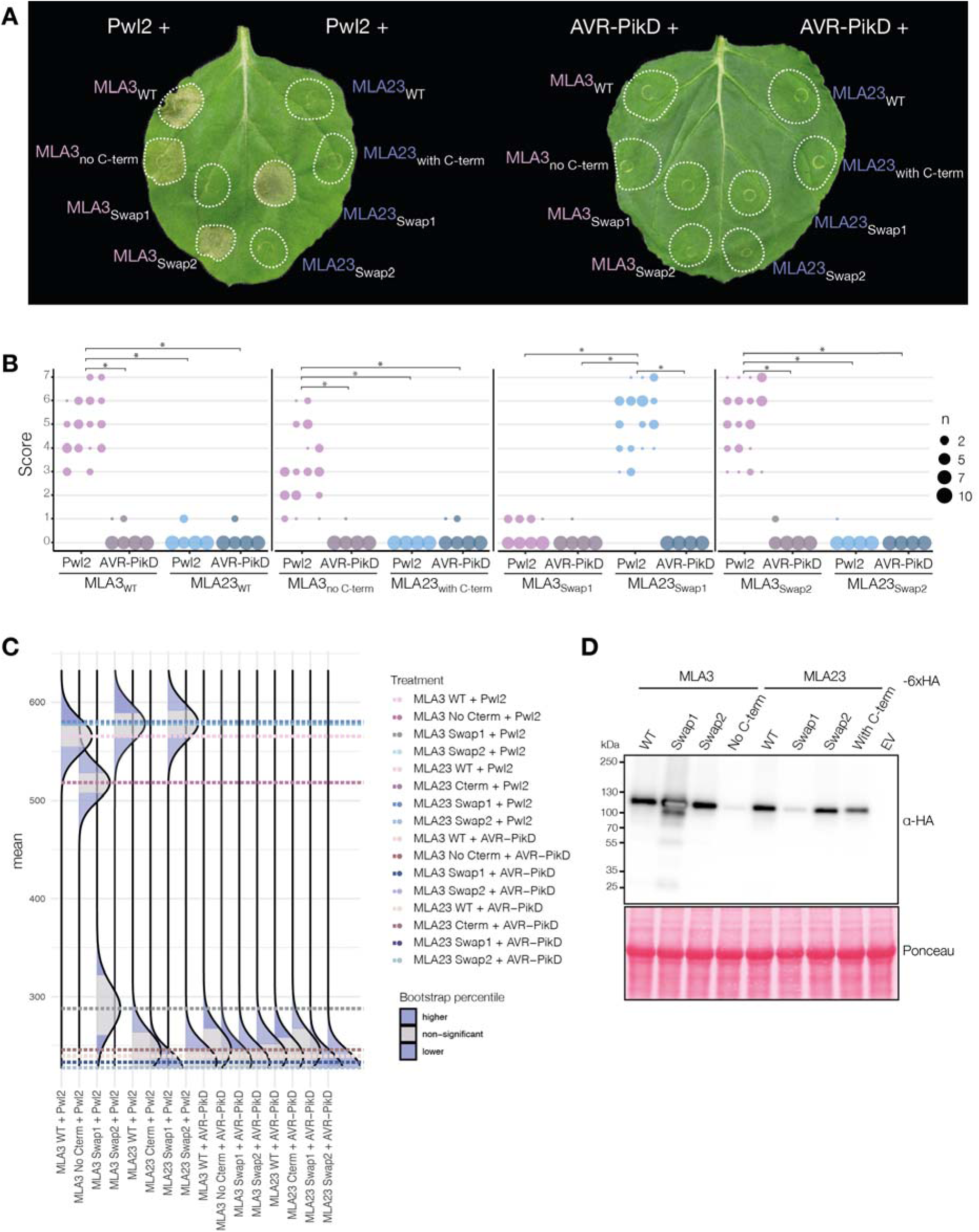
Introducing the unique MLA3 residues at the Pwl2 binding interface into MLA23 results in gain of Pwl2 recognition (extended Fig. 3F). (**A)** Representative photos of *N. benthamiana* leaves expressing the indicated chimeric constructs according to the swaps indicated in Fig. 3E. The *M. oryzae* effector AVR-PikD was used as negative control. Photos were taken three days post-agroinfiltration. (**B**) Cell death phenotypes from (A) were scored at three days post-agroinfiltration. The results are presented as a dot plot, where the size of a dot is proportional to the number of technical replicates with the same score within the same biological replicate. The experiments were independently repeated four times with at least 6 technical replicates; the three data point columns of each tested condition correspond to results from different biological replicates. Significant differences between conditions were calculated based on bootstrapping rank statistics and are indicated with an asterisk. (**C**) Statistical analysis of the cell death response shown in (B) performed with the estimation method from the besthr R library (*49*). Distribution of 1000 bootstrap sample rank means, with the areas in blue highlighting the 0.025 and 0.975 percentiles of the distribution. The grey area indicates the 95% confidence interval of the ranked mean. The difference of cell death response is considered statistically significant if the confidence interval of the ranked mean of a given treatment falls outside of the confidence interval of another condition (i.e. within or beyond the blue percentile of the mean distribution of another condition). (**D**) Anti-HA immunoblots of the C-terminally 6xHA tagged MLA chimeric proteins used in (A). Total protein extracts were prepared from leaves harvested 3 days after agroinfiltration. Protein loading was checked with Ponceau S solution.

**Fig. S11.**
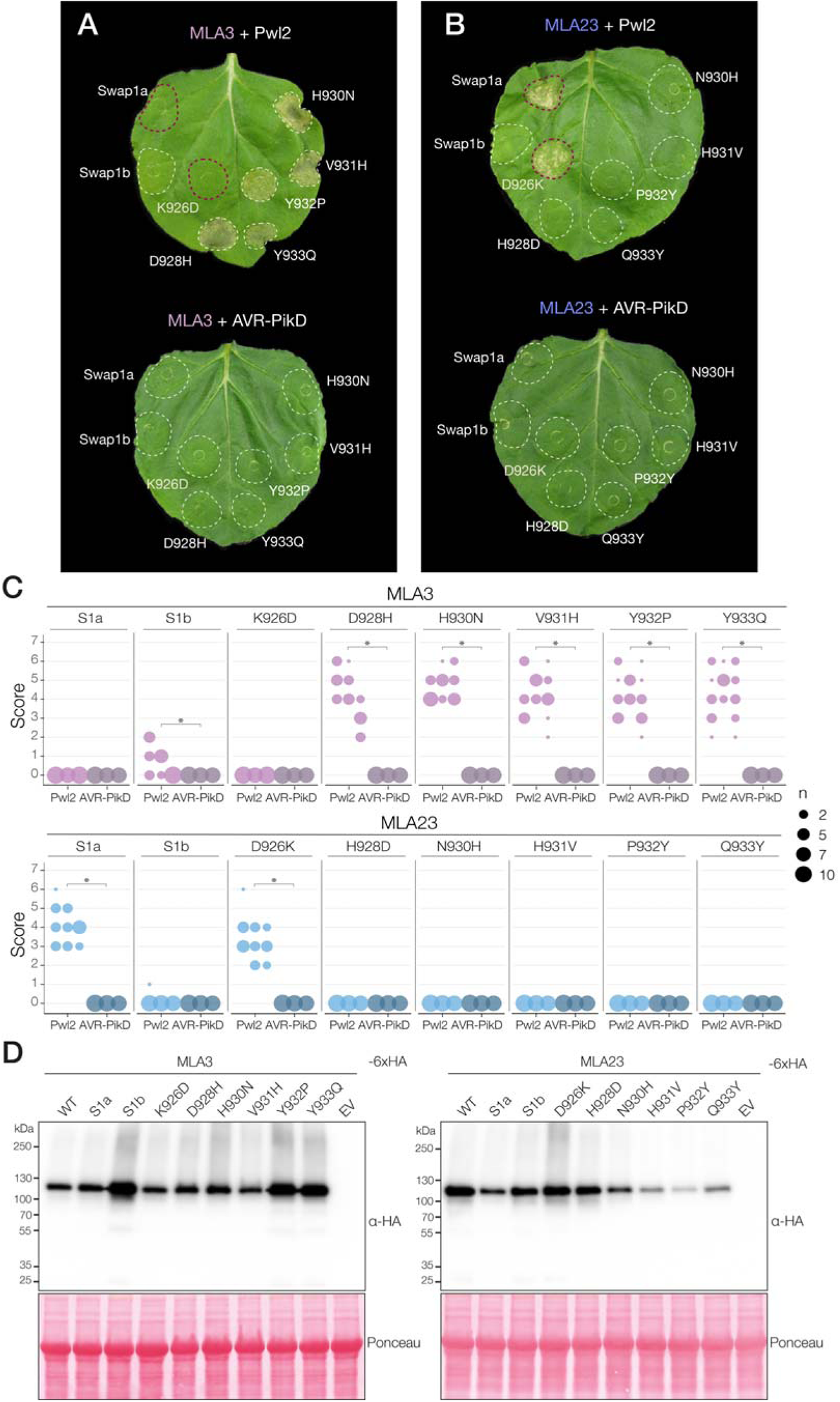
The D926K substitution in MLA23 is sufficient to confer Pwl2 recognition (extended Fig. 3G). (**A** and **B**) Representative photos of *N. benthamiana* leaves expressing the indicated chimeric proteins and reciprocal single amino acid substitutions in MLA3 (A) and MLA23 (B). The *M. oryzae* effector AVR-PikD was used as negative control. Red dashed circles highlight an exchanged cell death response between the MLA3 and MLA23 constructs. Photos were taken three days post-agroinfiltration. (**C**) Cell death phenotypes from (A) and (B) were scored at three days post-agroinfiltration. The results are presented as a dot plot, where the size of a dot is proportional to the number of technical replicates with the same score within the same biological replicate. The experiments were independently repeated three times with at least 6 technical replicates; the three data point columns of each tested condition correspond to results from different biological replicates. Significant differences between conditions were calculated based on bootstrapping rank statistics and are indicated with asterisk. (**D**) Anti-HA immunoblots of the C-terminally 6xHA tagged MLA proteins used in (A) and (B). Total protein extracts were prepared from leaves harvested 3 days after agroinfiltration. Protein loading was checked with Ponceau S solution.

**Fig. S12.**
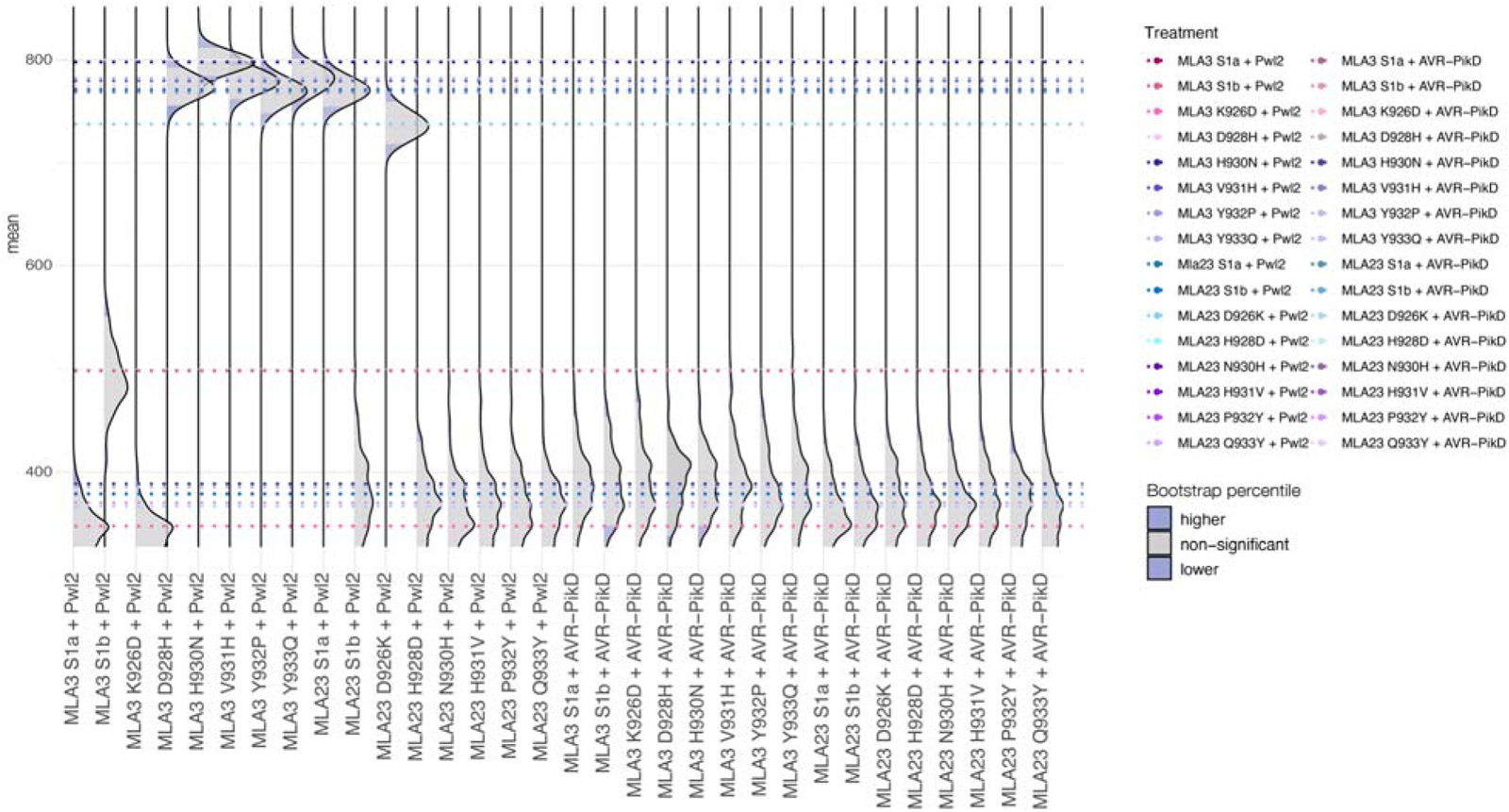
Statistical analysis of the quantification of the cell death phenotype of MLA3 and MLA23 Swap1a/b chimeras and single amino acid substitutions. Statistical analysis of the cell death response shown in (fig. S12C) performed with the estimation method from the besthr R library (*49*). Distribution of 1000 bootstrap sample rank means, with the areas in blue highlighting the 0.025 and 0.975 percentiles of the distribution. The grey area indicates the 95% confidence interval of the ranked mean. The difference of cell death response is considered statistically significant if the confidence interval of the ranked mean of a given treatment falls outside of the confidence interval of another condition (i.e. within or beyond the blue percentile of the mean distribution of another condition).

**Fig. S13.**
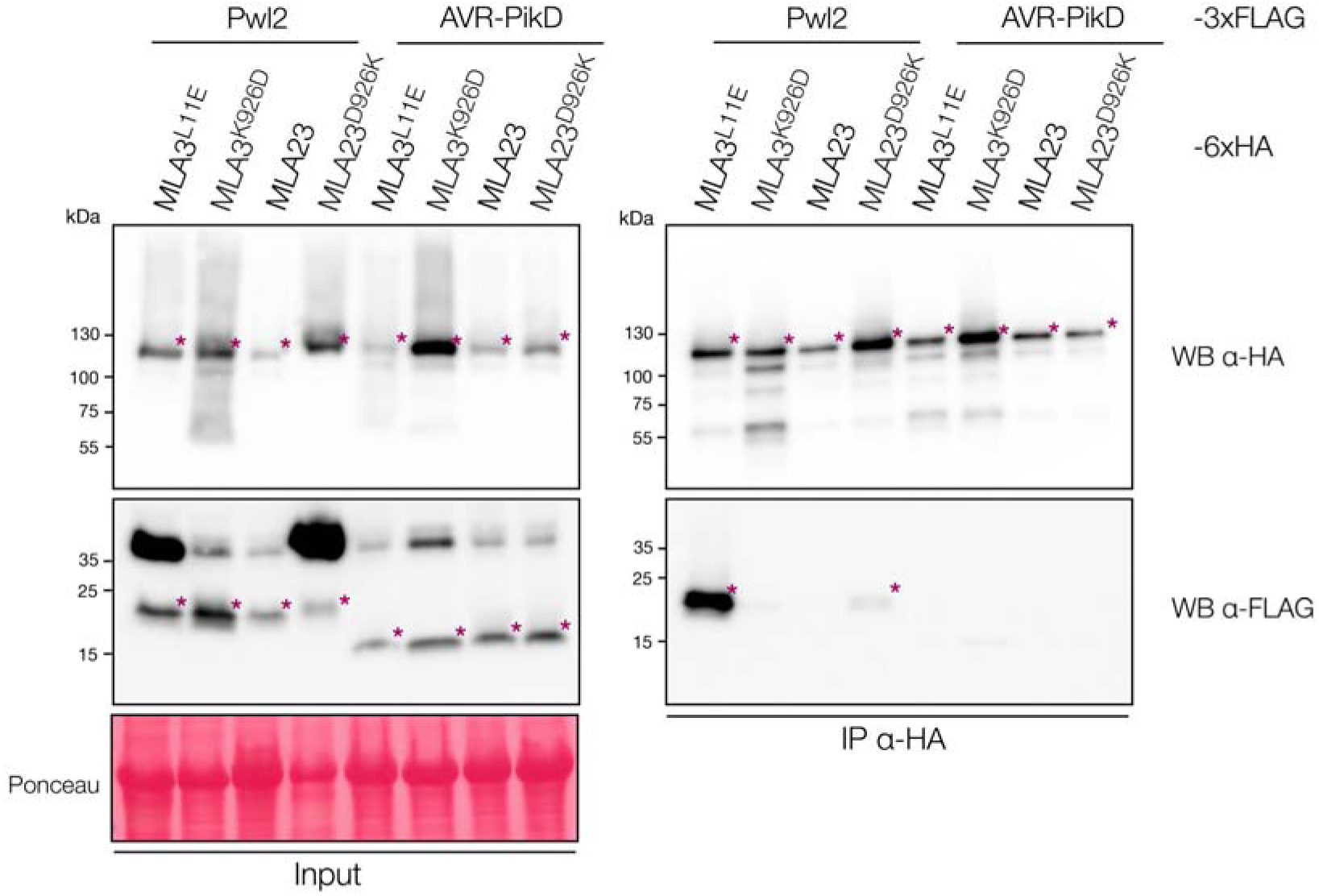
The D926K substitution in MLA23 is sufficient to gain Pwl2 binding. Coimmunoprecipitation of C-terminally tagged 6x-HA MLA3^L11E^, MLA3^K926D^, MLA23 or MLA23^D926K^ (∼115 kDa) with C-terminally tagged 3xFLAG Pwl2 (17.3 kDa) and AVR-PikD (14 kDa) in *N. benthamiana*. AVR-PikD was used as negative control. Total protein extracts were prepared from leaves harvested 3 days after agroinfiltration. Immunoprecipitation (IP) was performed with anti-HA agarose beads. Total protein (Input) and pulled-down fractions (IP) were detected with the antibodies labelled on the right. Asterisks show the expected protein band for each treatment. Protein loading was checked with Ponceau S solution. The experiment was independently repeated three times with similar results.

**Fig. S14.**
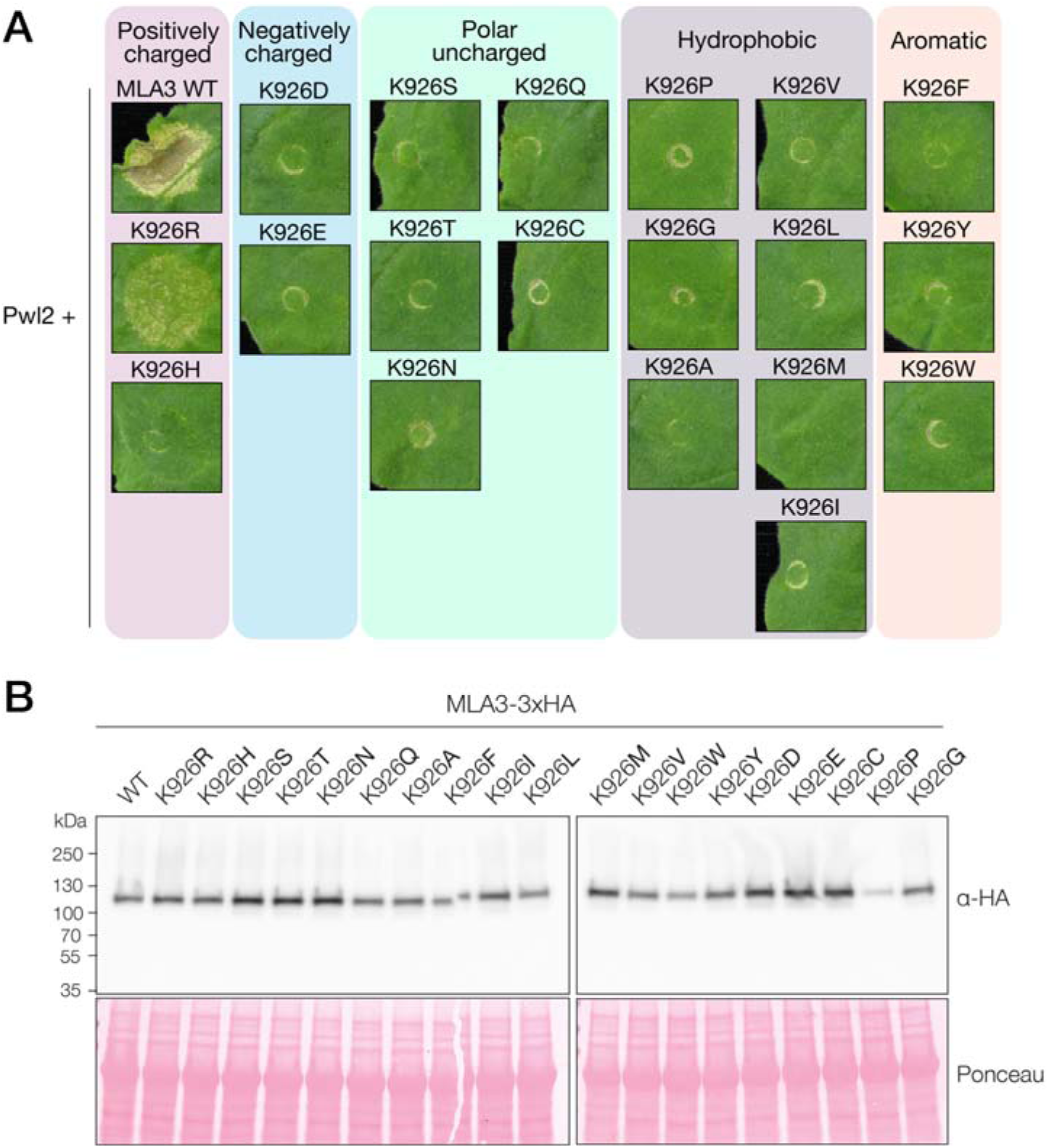
A positively charged residue is required in position 926 of MLA3 to confer Pwl2 recognition. (**A**) Representative *N. benthamiana* leaf panels showing HR response after co-expressing Pwl2 and the indicated MLA3 K926 substitutions. Photos were taken 3 days post agroinfiltration. The experiment was independently repeated two times with 6-8 technical replicates. **(B)** Anti-HA immunoblots of the C-terminally 6xHA tagged MLA3 variants (∼115 kDa) from (A) expressed in *N. benthamiana.* Total protein extracts were prepared from leaves harvested 3 days after agroinfiltration. Protein loading was checked with Ponceau S solution (Sigma).

**Fig. S15.**
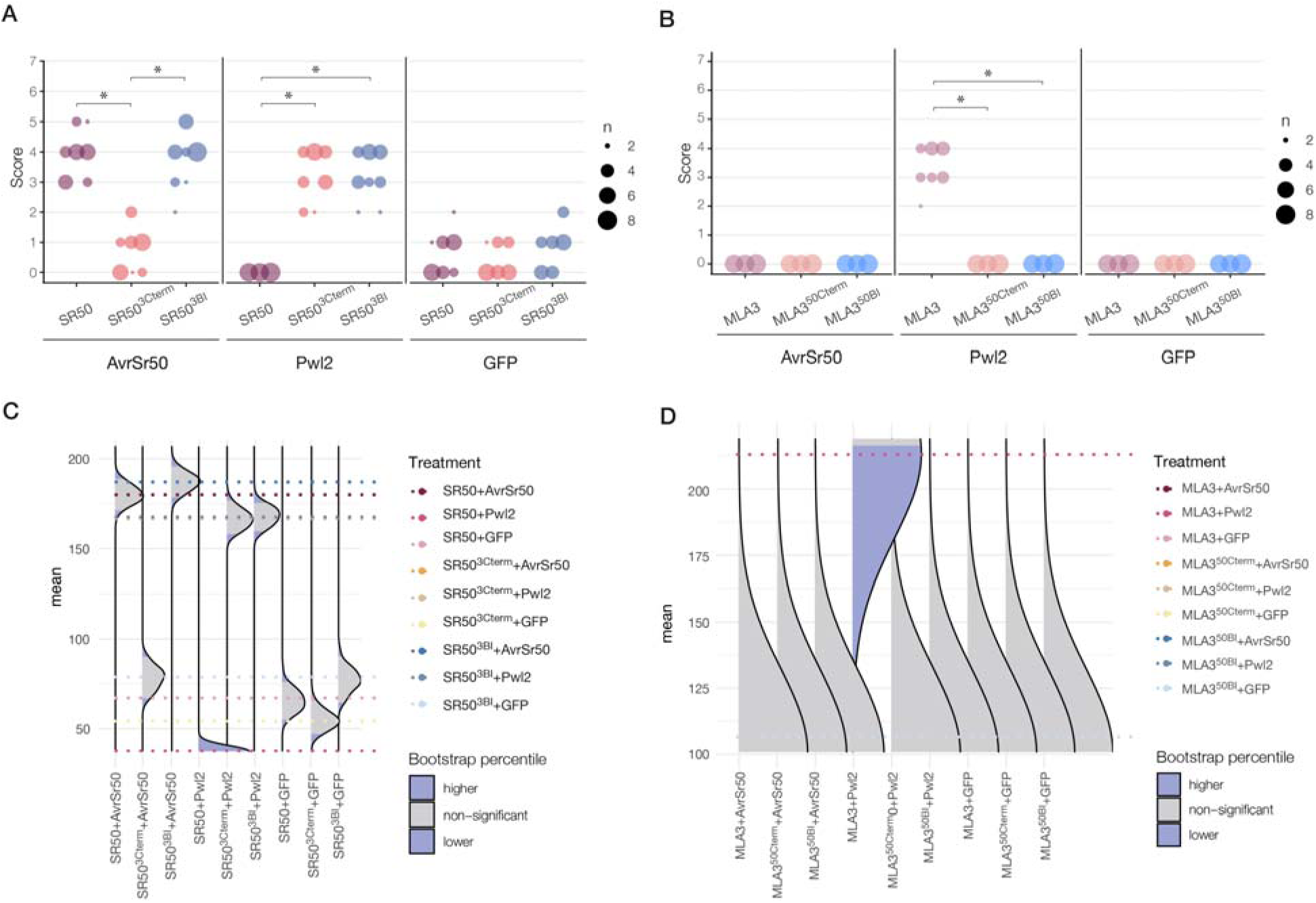
Cell death quantification of MLA3-SR50 chimeric proteins. (**A** and **B**) Cell death phenotypes from Fig. 4B**-C** were scored at three days post-agroinfiltration. Quantification of cell death triggered by SR50 (A) and MLA3 (B) chimeras when coexpressed with the indicated fungal effectors or GFP. The results are presented as a dot plot, where the size of a dot is proportional to the number of technical replicates with the same score within the same biological replicate. The experiments were independently repeated three times with at least 6 technical replicates; the three data point columns of each tested condition correspond to results from different biological replicates. Significant differences between conditions were calculated based on bootstrapping rank statistics and are indicated with asterisk. (**C** and **D**) Statistical analysis of the cell death phenotypes shown in (A) and (B) performed with the estimation method from the besthr R library (*49*) for the SR50 (C) and MLA3 (D) chimeras. Distribution of 1000 bootstrap sample rank means, with the areas in blue highlighting the 0.025 and 0.975 percentiles of the distribution. The grey area indicates the 95% confidence interval of the ranked mean. The difference of cell death response is considered statistically significant if the confidence interval of the ranked mean of a given treatment falls outside of the confidence interval of another condition (i.e. within or beyond the blue percentile of the mean distribution of another condition).

**Fig. S16.**
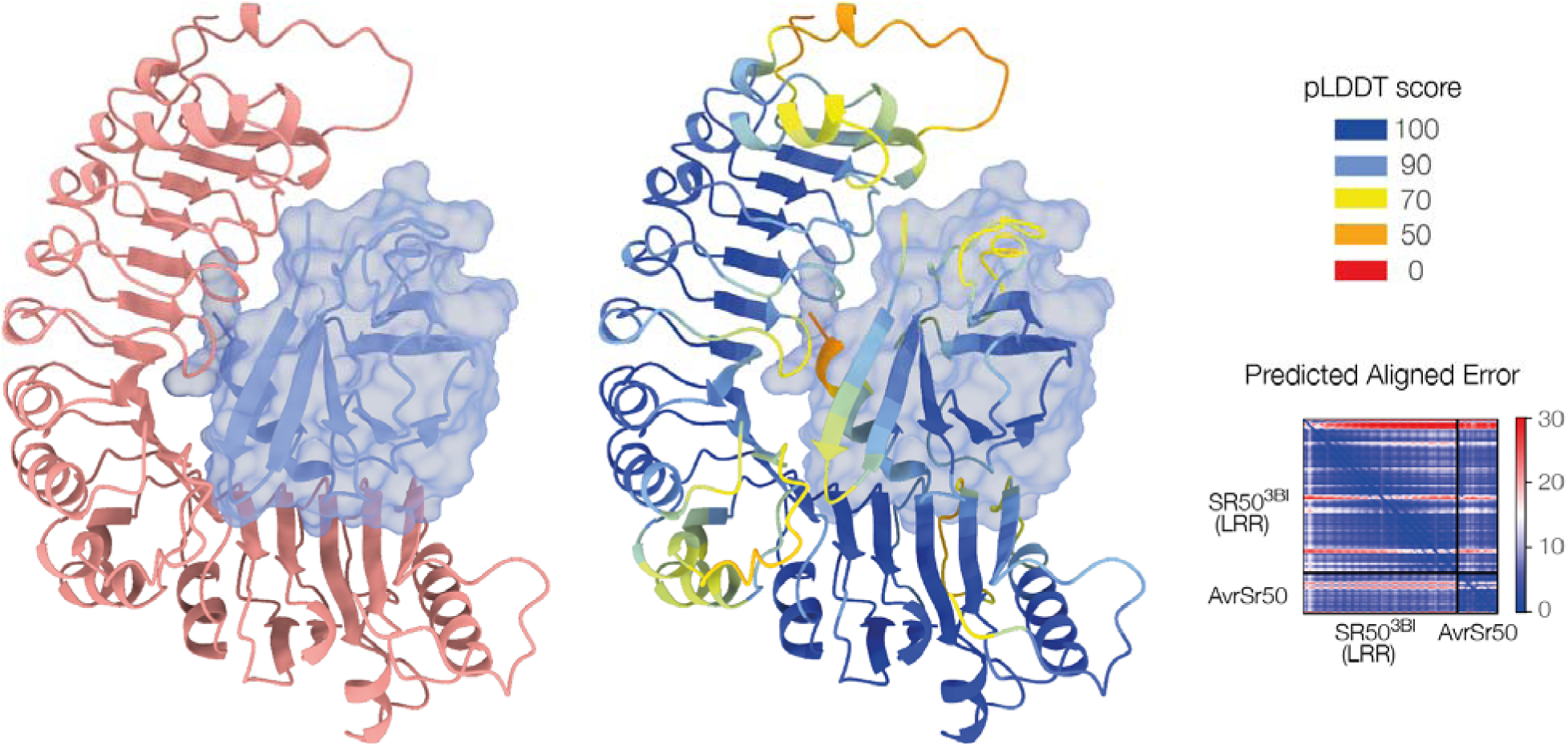
AlphaFold2 structure prediction of the LRR domain of the SR50^3BI^ chimera in complex with AvrSr50. The structure on the right is colored by pLDDT score and accompanied by the predicted aligned error plot.

**Table S1.**
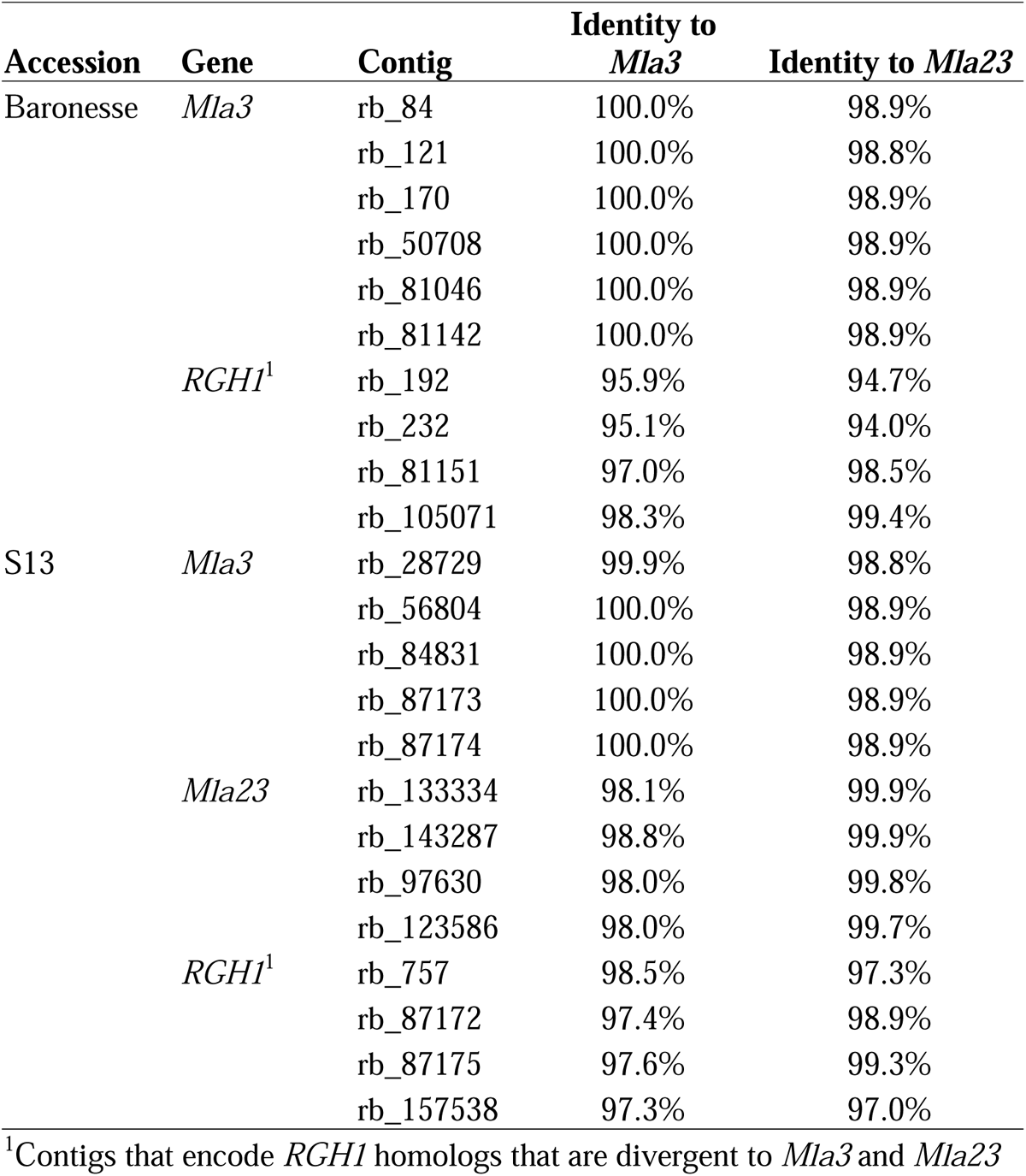
ONT *de novo* assembled cDNA contigs identity to the *Mla3* or *Mla23* alleles in the accessions Baronesse and S13.

**Table S2.**
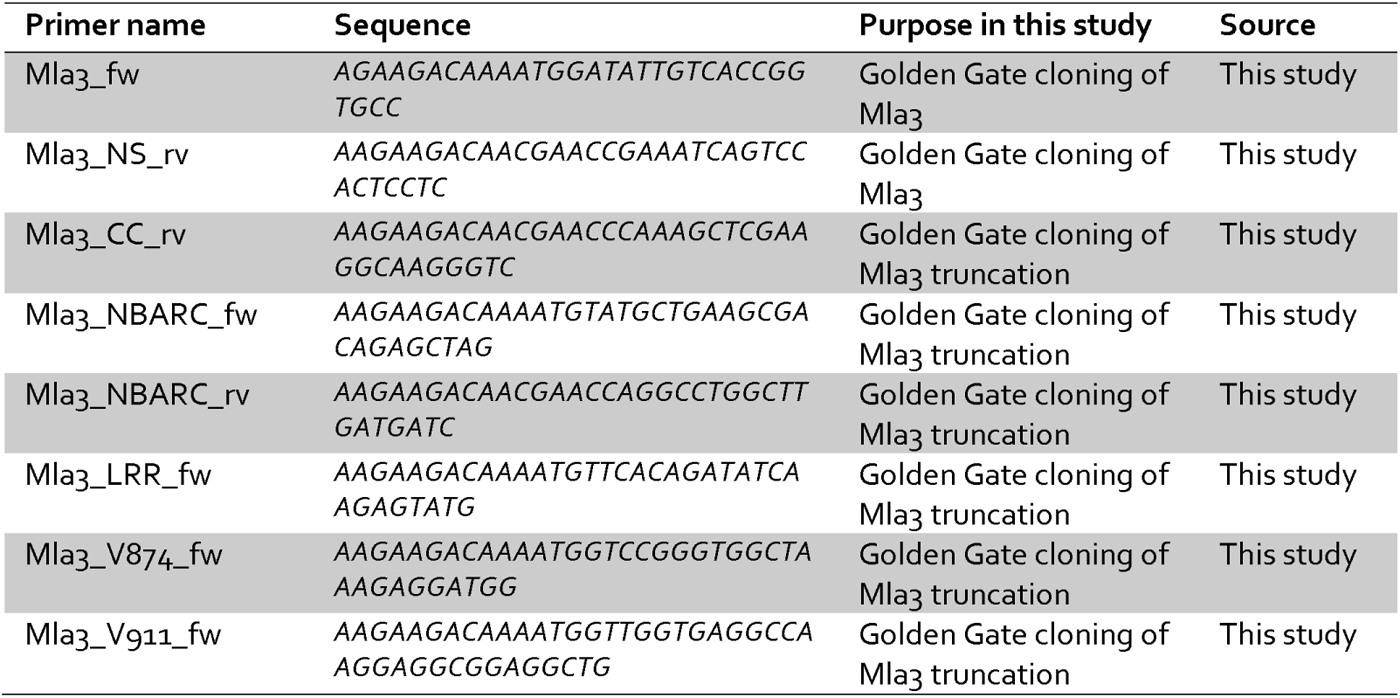
List of primers used to clone *Mla3* truncations.

**Table S3.**
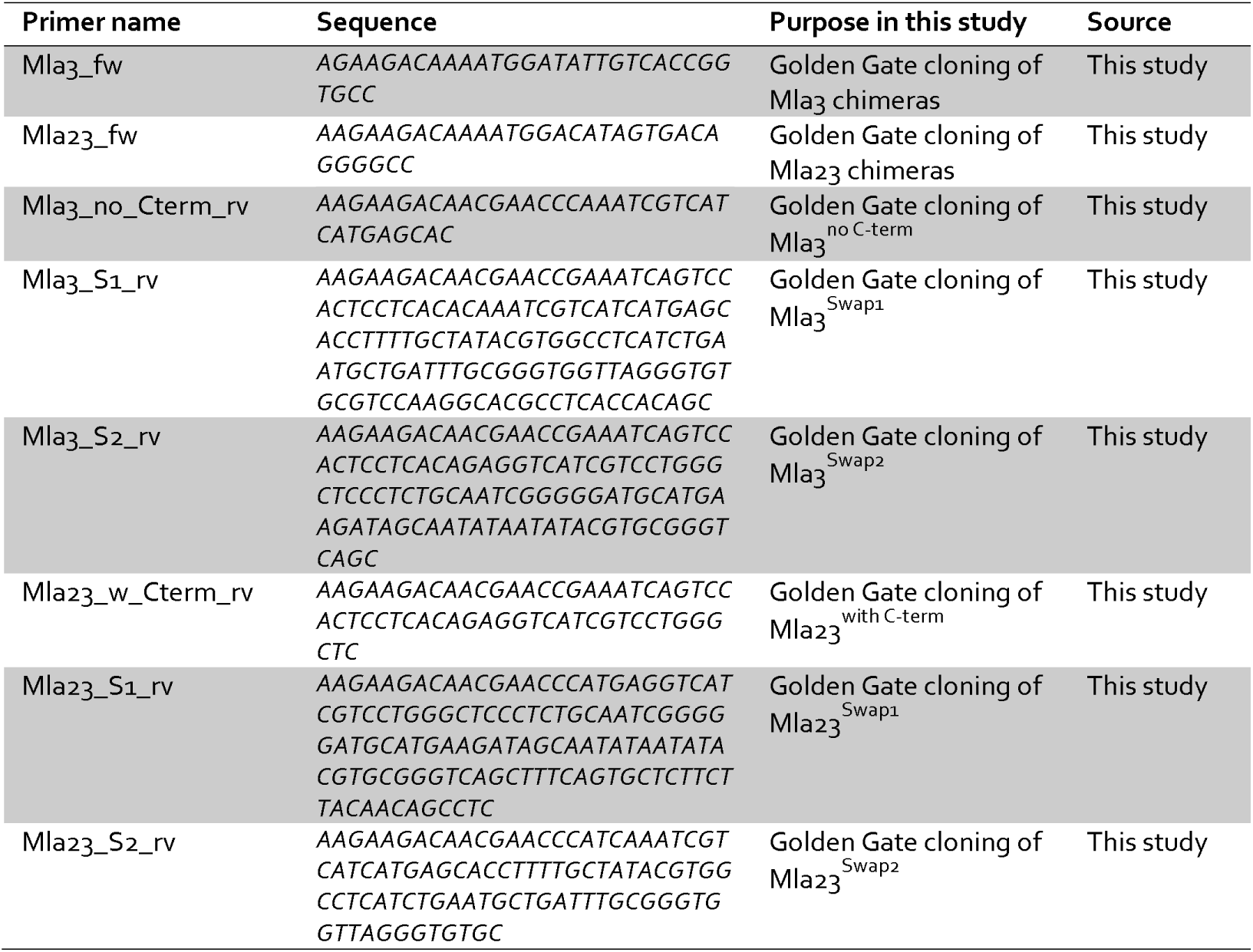
List of primers to clone *Mla3* and *Mla23* chimeras.

Table S4. Primers to clone *Mla3* and *Mla23* Swap1 mutants (as separate file).

Table S5. Oligos to generate *Mla3* K926 amino acid mutations (as separate file).

## References and Notes

1. J. D. G. Jones, R. E. Vance, J. L. Dangl, Intracellular innate immune surveillance devices in plants and animals. Science 354, aaf6395 (2016).

2. Z. Duxbury et al., Pathogen perception by NLRs in plants and animals: Parallel worlds. Bioessays 38, 769–781 (2016).

3. B. Sundaram, R. E. Tweedell, S. Prasanth Kumar, T.-D. Kanneganti, The NLR family of innate immune and cell death sensors. Immunity 57, 674–699 (2024).

4. L. A. Gao et al., Prokaryotic innate immunity through pattern recognition of conserved viral proteins. Science 377, eabm4096 (2022).

5. D. D. Leipe, E. V. Koonin, L. Aravind, STAND, a Class of P-Loop NTPases Including Animal and Plant Regulators of Programmed Cell Death: Multiple, Complex Domain Architectures, Unusual Phyletic Patterns, and Evolution by Horizontal Gene Transfer. J. Mol. Biol. 343, 1–28 (2004).

6. J. Kourelis, T. Sakai, H. Adachi, S. Kamoun, RefPlantNLR is a comprehensive collection of experimentally validated plant disease resistance proteins from the NLR family. PLoS Biol. 19, e3001124 (2021).

7. Z.-Q. Shao et al., Large-Scale Analyses of Angiosperm Nucleotide-Binding Site-Leucine-Rich Repeat Genes Reveal Three Anciently Diverged Classes with Distinct Evolutionary Patterns. Plant Physiol. 170, 2095–2109 (2016).

8. Z. Duxbury, C.-h. Wu, P. Ding, A Comparative Overview of the Intracellular Guardians of Plants and Animals: NLRs in Innate Immunity and Beyond. Annu. Rev. Plant Biol. 72, 155–184 (2021).

9. R. A. L. van der Hoorn, S. Kamoun, From Guard to Decoy: A new model for perception of plant pathogen effectors. Plant Cell 20, 2009–2017 (2008).

10. J. Kourelis, R. A. L. van der Hoorn, Defended to the Nines: 25 Years of Resistance Gene Cloning Identifies Nine Mechanisms for R Protein Function. The Plant Cell 30, 285–299 (2018).

11. S. Cesari, M. Bernoux, P. Moncuquet, T. Kroj, P. Dodds, A novel conserved mechanism for plant NLR protein pairs: the ‘integrated decoy’ hypothesis. Frontiers in Plant Science 5, (2014).

12. S. Cesari, Multiple strategies for pathogen perception by plant immune receptors. New Phytol. 219, 17–24 (2018).

13. M. P. Contreras, D. Lüdke, H. Pai, A. Toghani, S. Kamoun, NLR receptors in plant immunity: making sense of the alphabet soup. EMBO reports 24, e57495 (2023).

14. S. Seeholzer et al., Diversity at the Mla Powdery Mildew Resistance Locus from Cultivated Barley Reveals Sites of Positive Selection. Mol. Plant-Microbe Interact. 23, 497–509 (2010).

15. H. J. Brabham et al., Barley MLA3 recognizes the host-specificity effector Pwl2 from Magnaporthe oryzae. The Plant Cell 36, 447–470 (2024).

16. J. A. Sweigard et al., Identification, cloning, and characterization of PWL2, a gene for host species specificity in the rice blast fungus. The Plant Cell 7, 1221–1233 (1995).

17. V. Were et al., The blast effector Pwl2 is a virulence factor that modifies the cellular localisation of host protein HIPP43 to suppress immunity. bioRxiv, 2024.2001.2020.576406 (2024).

18. S. M. Latorre et al., Differential loss of effector genes in three recently expanded pandemic clonal lineages of the rice blast fungus. BMC Biol. 18, 88 (2020).

19. A. Förderer, D. Yu, E. Li, J. Chai, Resistosomes at the interface of pathogens and plants. Curr. Opin. Plant Biol. 67, 102212 (2022).

20. A. Förderer, J. Kourelis, NLR immune receptors: structure and function in plant disease resistance. Biochem. Soc. Trans. 51, 1473–1483 (2023).

21. H. Adachi et al., An N-terminal motif in NLR immune receptors is functionally conserved across distantly related plant species. eLife 8, e49956 (2019).

22. R. Martin et al., Structure of the activated ROQ1 resistosome directly recognizing the pathogen effector XopQ. Science 370, eabd9993 (2020).

23. S. Ma et al., Direct pathogen-induced assembly of an NLR immune receptor complex to form a holoenzyme. Science 370, eabe3069 (2020).

24. M. Ravensdale, et al., Intramolecular Interaction Influences Binding of the Flax L5 and L6 Resistance Proteins to their AvrL567 Ligands. PLoS Path. 8, e1003004 (2012).

25. A. Förderer et al., A wheat resistosome defines common principles of immune receptor channels. Nature 610, 532–539 (2022).

26. Y.-B. Zhao et al., Pathogen effector AvrSr35 triggers Sr35 resistosome assembly via a direct recognition mechanism. Science Advances 8, eabq5108 (2022).

27. J. Jumper et al., Highly accurate protein structure prediction with AlphaFold. Nature 596, 583–589 (2021).

28. R. Evans et al. Protein complex prediction with AlphaFold-Multimer. bioRxiv, 2021.2010.2004.463034 (2021).

29. K. de Guillen et al., Structure Analysis Uncovers a Highly Diverse but Structurally Conserved Effector Family in Phytopathogenic Fungi. PLoS Path. 11, e1005228 (2015).

30. R. Zdrzałek et al., Bioengineering a plant NLR immune receptor with a robust binding interface toward a conserved fungal pathogen effector. Proceedings of the National Academy of Sciences 121, e2402872121 (2024).

31. E. Krissinel, K. Henrick, Inference of Macromolecular Assemblies from Crystalline State. J. Mol. Biol. 372, 774–797 (2007).

32. E. Krissinel, Crystal contacts as nature’s docking solutions. Journal of Computational Chemistry 31, 133–143 (2010).

33. J. Chen et al., Loss of AvrSr50 by somatic exchange in stem rust leads to virulence for Sr50 resistance in wheat. Science 358, 1607–1610 (2017).

34. R. Mago et al., The wheat Sr50 gene reveals rich diversity at a cereal disease resistance locus. Nature Plants 1, 15186 (2015).

35. J. Tamborski, K. Seong, F. Liu, B. J. Staskawicz, Ksenia V. Krasileva, Altering Specificity and Autoactivity of Plant Immune Receptors Sr33 and Sr50 Via a Rational Engineering Approach. Molecular Plant-Microbe Interactions® 36, 434–446 (2023).

36. D. Ortiz et al., The stem rust effector protein AvrSr50 escapes Sr50 recognition by a substitution in a single surface-exposed residue. New Phytol. 234, 592–606 (2022).

37. N. C. Elde, H. S. Malik, The evolutionary conundrum of pathogen mimicry. Nature Reviews Microbiology 7, 787–797 (2009).

38. H. Li et al., Pathogen protein modularity enables elaborate mimicry of a host phosphatase. Cell 186, 3196–3207.e3117 (2023).

39. P. Ronald, A. Joe, Molecular mimicry modulates plant host responses to pathogens. Ann. Bot. 121, 17–23 (2017).

40. D. Wu, L. Wang, Y. Zhang, L. Bai, F. Yu, Emerging roles of pathogen-secreted host mimics in plant disease development. Trends Parasitol. 37, 1082–1095 (2021).

41. S. Mondino, S. Schmidt, C. Buchrieser, Molecular Mimicry: a Paradigm of Host-Microbe Coevolution Illustrated by Legionella. mBio 11, 10.1128/mbio.01201-01220 (2020).

42. C. E. Stebbins, J. E. Galán, Structural mimicry in bacterial virulence. Nature 412, 701–705 (2001).

43. C. E. Stebbins, J. E. Galán, Modulation of Host Signaling by a Bacterial Mimic: Structure of the *Salmonella* Effector SptP Bound to Rac1. Mol. Cell 6, 1449–1460 (2000).

44. P. K. Singh et al., Salmonella SipA mimics a cognate SNARE for host Syntaxin8 to promote fusion with early endosomes. J. Cell Biol. 217, 4199–4214 (2018).

45. K. Seong, K. V. Krasileva, Prediction of effector protein structures from fungal phytopathogens enables evolutionary analyses. Nature Microbiology 8, 174–187 (2023).

46. Y. Cao et al., Structural polymorphisms within a common powdery mildew effector scaffold as a driver of coevolution with cereal immune receptors. Proceedings of the National Academy of Sciences 120, e2307604120 (2023).

47. S. Marillonnet, R. Grützner, Synthetic DNA Assembly Using Golden Gate Cloning and the Hierarchical Modular Cloning Pipeline. Current Protocols in Molecular Biology 130, e115 (2020).

48. A. Maqbool et al., Structural basis of pathogen recognition by an integrated HMA domain in a plant NLR immune receptor. eLife 4, e08709 (2015).

49. D. MacLean. (Zenodo, 2019).

